# The outer membrane lipoprotein NlpI nucleates hydrolases within peptidoglycan multi-enzyme complexes in *Escherichia coli*

**DOI:** 10.1101/609503

**Authors:** Manuel Banzhaf, Hamish C. L. Yau, Jolanda Verheul, Adam Lodge, George Kritikos, André Mateus, Ann Kristin Hov, Frank Stein, Morgane Wartel, Manuel Pazos, Alexandra S. Solovyova, Mikhail M Savitski, Tanneke den Blaauwen, Athanasios Typas, Waldemar Vollmer

## Abstract

The peptidoglycan (PG) sacculus provides bacteria with the mechanical strength to maintain cell shape and resist osmotic stress. Enlargement of the mesh-like sacculus requires the combined activity of PG synthases and hydrolases. In *Escherichia coli*, the activity of the two bifunctional PG synthases is driven by lipoproteins anchored in the outer membrane. However, the regulation of PG hydrolases is less well understood, with only regulators for PG amidases having been described. Here, we identify the lipoprotein NlpI as a general adaptor protein for PG hydrolases. NlpI binds to different classes of hydrolases and can specifically form multimeric complexes with various PG endopeptidases. In addition, NlpI seems to contribute both to PG elongation and cell division biosynthetic complexes based on its localization and genetic interactions. In line with such a role, we reconstitute PG multi-enzyme complexes containing NlpI, the PG synthesis regulator LpoA, its cognate bifunctional synthase, PBP1A, and different endopeptidases. Our results indicate that PG regulators and adaptors are part of PG biosynthetic multi-enzyme complexes, regulating and potentially coordinating the spatiotemporal action of PG synthases and hydrolases.

**Significance:** The activity of PG hydrolases may cause lysis of the bacterial cell if left unregulated. Hence, the cell must have ways of regulating and coordinating their activities. Our current understanding of how this occurs is incomplete. In this work, we present the outer membrane (OM) anchored lipoprotein, NlpI, as a scaffold of peptidoglycan hydrolases. We propose that NlpI facilitates the formation of multi-enzyme complexes and that, along with other regulators, it coordinates a safe enlargement and separation of the PG layer in *E. coli*.

## Introduction

Peptidoglycan (PG) provides bacteria with the mechanical strength to maintain cell shape and resist osmotic stresses. The PG layer or sacculus is a mesh-like structure composed of glycan chains connected by peptides and surrounds the cytoplasmic membrane (CM) (Silhavy *et al.*, 2010; Vollmer *et al.*, 2008a). Given the internal turgor of the cells, PG layer growth requires the coordinated action of synthases and hydrolases to enlarge the sacculus without rupture. This important task is executed by large protein complexes, the elongasome and the divisome, which recruit PG enzymes together with regulators, cytoskeletal, morphogenesis and other structural proteins (den Blaauwen *et al.*, 2017; Typas *et al.*, 2012; Typas & Sourjik, 2015). It has been previously hypothesized that the formation of these complexes enables the cell to coordinate and regulate the activities of various synthetic and hydrolytic PG enzymes in a spatiotemporal manner (Höltje, 1993). Within these complexes, the key bifunctional penicillin-binding protein (PBP) PG synthases are activated by cognate outer membrane (OM)-anchored lipoproteins (Dorr *et al.*, 2014; More *et al.*, 2019; Paradis-Bleau *et al.*, 2010; Typas *et al.*, 2010; Typas *et al.*, 2012) and coordinate their action with other constriction related protein complexes (Gray *et al.*, 2015). However, with the exception of the amidases (Peters *et al.*, 2013; Tsang *et al.*, 2017; Uehara *et al.*, 2010; Yang *et al.*, 2012), it is less clear how Gram-negative bacteria control the activities of their wide repertoire of hydrolases – i.e. the endopeptidases (EPases), carboxypeptidases (CPases) and lytic transglycosylases (LTases).

NlpI is an OM-anchored lipoprotein predicted to be involved in cell division and responsible for targeting the PG EPase MepS for proteolytic degradation (Ohara *et al.*, 1999; Singh *et al.*, 2015). Deletion of *nlpI* causes cell filamentation at elevated temperature (42°C) or low osmolarity, whilst overexpresing NlpI results in the formation of prolate spheroids (Ohara *et al.*, 1999). Deletion of *nlpI* has further implications on the stability of the OM as it leads to hypervesiculation in a manner that is dependent on the activity of two EPases; PBP4 in stationary phase and MepS in exponential phase (Schwechheimer *et al.*, 2015). This phenotype is suppressed by a deletion of *mepS* (Schwechheimer *et al.*, 2015). Many of its pleiotropic effects may be due to the ability of NlpI to target the EPase MepS for proteolytic degradation by forming a complex with the tail-specific protease Prc (Su *et al.*, 2017). NlpI and MepS both interact with Prc, but whilst MepS is digested, only 12 C-terminal amino acids of NlpI are removed (Singh *et al.*, 2015). In the absence of NlpI, the half-life of MepS increases from ~2 min to ~45 min. Further in the Δ*nlpI* mutant, uncontrolled levels of MepS have been shown to impair cell growth on low osmolarity medium and lead to the formation of long filaments (Singh *et al.*, 2012; Singh *et al.*, 2015).

NlpI forms a homodimer (Wilson *et al.*, 2005) with the 33 kDa monomers having their OM-binding N-termini in close proximity. Each monomer consists of 14 α-helices forming 4 canonical but distinct tetratricopeptide repeat (TPR)-like domains, which are characteristic for protein interacting modules. A putative binding cleft is formed from the curvature of the helices on each monomer, which would be available for protein-protein interactions (Das *et al.*, 1998; Wilson *et al.*, 2005). It is hence possible that NlpI acts as a scaffold for the formation of protein complexes. In this study, we provide evidence that the primary cellular function of NlpI is to nucleate hydrolases within PG multi-enzyme complexes in *E. coli*.

## Results

### Deletion of NlpI alters abundance and thermostability of envelope biogenesis proteins

Deletion of *nlpI* causes several pleiotropic phenotypes. To link the observed phenotypes to changes in protein abundance and activity, we compared an *nlpI* knockout strain (Δ*nlpI*) to wildtype *E. coli* using two-dimensional Thermal Proteome Profiling (2D-TPP) (Mateus *et al.*, 2018; Savitski *et al.*, 2014). In TPP, both protein abundance and thermostability can be measured. The latter depends on the intrinsic physical properties of the protein and on external factors that stabilize its fold; such as protein-protein and protein-ligand interactions.

Numerous proteins changed abundance and thermostability in the Δ*nlpI* cells (supplementary table 1 and 2). In agreement with its periplasmic location and link to envelope integrity (Schwechheimer *et al.*, 2015), deletion of *nlpI* resulted in changes in abundance and thermostability of major envelope components, including outer membrane (OMPs), the β-barrel assembly machinery (BAM) (Noinaj *et al.*, 2017) and the Tol-Pal complex (Egan, 2018) (Fig. 1a and 1b). As expected, both MepS abundance and thermostability were significantly elevated in Δ*nlpI* cells, since in the absence of NlpI, MepS is not targeted for degradation by Prc (Singh *et al.*, 2015) (Fig. 1a and 1b). We also observed that other PG biogenesis proteins were affected. These included several PG hydrolases (MepS, PBP5, PBP6a, MltA, MltG, LdtB, LdtF) and PG synthases (PBP1A, PBP1B), which increased in abundance in the Δ*nlpI* cells (Fig. 1a). A number of them also decreased in thermostability, with LTases (MltA, MltC, MltE), the LD-transpeptidase LdtF and the PG synthases and their regulators (PBP1B, LpoA, LpoB) showing a strong effect (Fig. 1b). In contrast, all amidases (AmiA, AmiB and AmiC) decreased in abundance (Fig. 1a). Moreover, the amidase regulator NlpD (which binds to AmiC and controls its activity) (Uehara *et al.*, 2010) and the YraP protein, which was recently implicated in the activation of NlpD, were strongly destabilised (Fig. 1b) (Tsang *et al.*, 2017).

**Figure 1.**
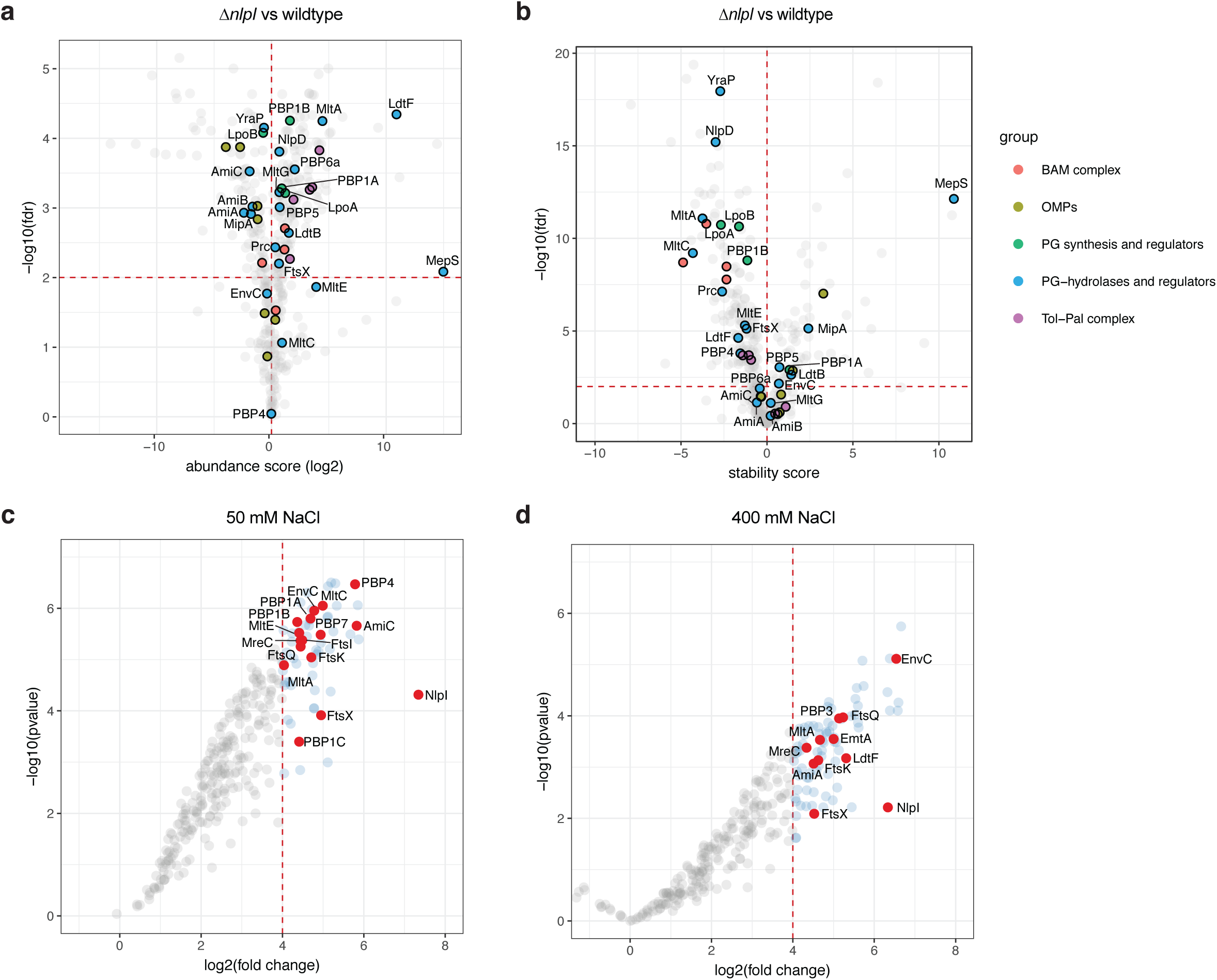
*In vivo* and *in vitro* proteomics-based assays link NlpI to several classes of PG hyrdolases. **a-b.** Wild-type and Δ*nlpI* cells were heated at a range of temperatures and the soluble components were labelled by TMT, combined and quantified by LC-MS, using the published 2D-TPP protocol (Mateus *et al.*, 2018). Shown are volcano plots of two replicates depicting changes in: protein abundance (a) and thermostability (b). A local FDR<0.01 was set as a threshold for significance. Highlighted proteins: outer membrane porins (OMPs, light green), β-barrel assembly machinery (BAMs, red), PG synthases and regulators (green), PG hydrolases and regulators (blue) and the Tol-Pal complex (violet). All other proteins were colored grey and not labelled to increase the plot clarity. Full results can be found in supplementary tables 1 and 2. **c-d**. Affinity chromatography with immobilised NlpI. Membrane extracts from *E. coli* were incubated in low and high salt binding conditions (50 and 400 mM NaCl, respectively), and then eluted with 1 M NaCl to identify possible interaction partners by label-free LC-MS analysis. Here plotted the log2 fold enrichment of proteins when compared to those eluted from a parallel empty column control, versus the log10 p-value, in low (4 replicates) (**c**) and high (2 replicates) (**d**) salt. Highlighted points are all interactions with PG enzymes and their regulators, as well as members of the divisome. All other proteins were colored grey and not labelled to increase the plot clarity; many were non-physiological interactions with abundant cytoplasmic proteins. Full results can be found in supplementary tables 3 and 4.

To rule out that all the observed changes are an indirect result of the higher abundance of MepS in the Δ*nlpI* strain, we repeated the 2D-TPP with a Δ*nlpI*Δ*mepS* strain (Fig. S1a and S1b). Several of the changes observed in the Δ*nlpI* cells remained in the Δ*nlpI*Δ*mepS* background (Fig. S1a and S1b), including the destabilization of many cell wall enzymes and regulators. We also compared directly the 2D-TPP profiles of Δ*nlpI* and Δ*nlpI*Δ*mepS* strains (Fig. S1c and S1d), with the major difference between both proteomes being that some OMPs were more stable in Δ*nlpI* cells. Importantly, the stability changes occurring for PG enzymes were not observed in this comparison, indicating that they occur independently of MepS levels. Altogether, these results provide the first evidence that NlpI affects PG biogenesis beyond the known interaction with the EPase MepS.

### NlpI pulls down several classes of PG hydrolases and multiple divisome proteins

The decrease in thermostability of several PG biogenesis proteins in Δ*nlpI* cells raised the possibility that NlpI may interact with these proteins. To investigate this further, we applied detergent solubilized *E. coli* membrane proteins to immobilised NlpI to identify potential interaction partners. Affinity chromatography was performed both in low salt binding conditions (50 mM) to pull down larger PG multi-enzyme complexes, and in high salt binding conditions (400 mM) to identify stronger, salt-resistant and possibly direct binding partners. As a control, we used a column containing Tris-coupled sepharose beads, and compared elution fractions with label-free mass spectrometry (supplementary table 3 and 4). To investigate relevant and direct NlpI interaction partners we focused on proteins located in the periplasmic space and highlighted known PG biogenesis proteins (Fig. 1c and 1d).

In low salt binding conditions NlpI retained several envelope biogenesis proteins, such as the PG synthases PBP1A, PBP1B, PBP1C, the divisome proteins EnvC, PBP3, FtsK, FtsQ and FtsX, the LTases MltA and MltC, the amidase AmiC and the EPases PBP4 and PBP7, amongst others (Fig. 1c). This shows that NlpI is able to pull-down full or partial PG-synthase complexes. When challenged in high salt binding conditions many of the aforementioned interactions were lost. Immobilized NlpI still retained the divisome proteins PBP3, FtsK, FtsQ and FtsX, the amidase AmiA and its regulator EnvC, and the LTase MltA at 400 mM NaCl, suggesting strong, salt-resistant interactions (Fig. 1d). Thus, in accordance with its phenotypes, NlpI interacts with the divisome and in accordance with the 2D-TPP results it may directly interact with several PG hydrolases (amidases, EPases and LTases).

The *in vivo* proteomics of Δ*nlpI* and the subsequent affinity chromatography revealed strong links of NlpI to several classes of PG hydrolases, PG synthases and divisome proteins. To investigate whether NlpI has a broader role in regulating EPases beyond MepS (Singh *et al.*, 2015), we next focused on characterizing the interactions of NlpI with EPases and PG synthases in more detail.

### NlpI dimerizes and interacts with several endopeptidases

To confirm the observed interactions between NlpI and EPases, we performed various biochemical assays. Firstly, we determined that NlpI is predominantly a homodimer using analytical ultracentrifugation (AUC). The experimentally determined sedimentation coefficient was 4.16 S which is close to the calculated sedimentation coefficient of 4.52, based on the crystal structure of the NlpI dimer (1XNF.pdb; Wilson *et al.*, 2005) (Fig. S2a). We measured the apparent dissociation constant (K_D_) for the NlpI dimer as 126 ± 9 nM by microscale thermophoresis (MST): titrating a fluorescently-labelled NlpI (fl-NlpI) against a serial dilution of unlabelled NlpI (Fig. 2a and S2b). The formation of a dimer by NlpI in solution is consistent with previous work (Su *et al.*, 2017). We next tested the specificity of a previously reported interaction between NlpI and the EPase MepS, using MST (Singh *et al.*, 2015). We found that NlpI and MepS interacted directly, with an apparent K_D_ of 145 ± 52 nM (Fig. 2a and 2b). NlpI interacted with MepM and PBP4 with similar apparent K_D_’s of 152 ± 42 nM and 177 ± 49 nM, respectively (Fig. 2a and Fig S2b). Assaying for an interaction between NlpI and PBP7 by MST revealed a more complex binding curve, which could only be fit assuming a Hill coefficient of ~ 3 (Fig. S2b). This resulted in an apparent EC_50_ value of 422 ± 25 nM and suggested an element of positive co-operativity in the NlpI-PBP7 binding.

**Figure 2.**
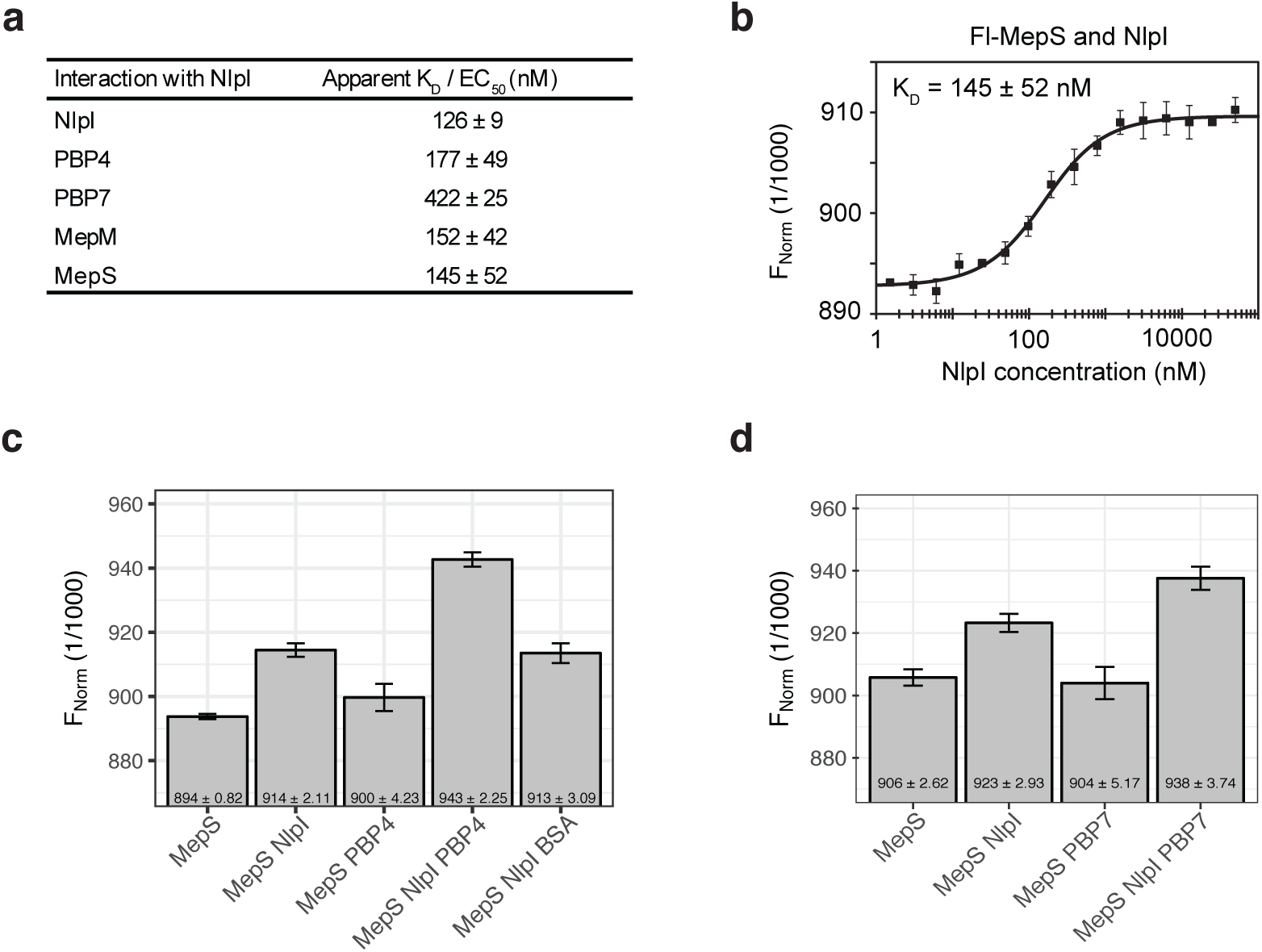
NlpI interacts with several EPases and is able to form trimeric complexes with them. **a.** Dissociation constants for interactions between NlpI with MepM, MepS, PBP4, PBP7 as determined by Microscale thermophoresis (MST). The values are mean ± standard deviation of three independent experiments. The corresponding MST binding curves are shown in Fig S2b. **b.** MepS-NlpI interaction by MST. Fl, fluorescently labelled; FNorm, normalized fluorescence. **c-d.** NlpI has different binding sites for MepS and PBP4, and MepS and PBP7 as shown by the ability of labelled MepS to bind a preformed NlpI-PBP4 complex (**c**) and NlpI-PBP7 complex (**d**) by a fixed concentration MST assay. Values are mean ± standard deviation of 3-6 independent experiments.

We also tested the interactions between NlpI and EPases (MepM, MepS, PBP4 and PBP7) by Ni^2+^-NTA pull down assays and confirmed the interactions found by MST (Fig. S3a). We could not detect an interaction between NlpI and the carboxypeptidase PBP5 or the LTase Slt, suggesting that NlpI does not interact with all hydrolases in general (Fig. S3a). Using a combination of MST and Ni^2+^-NTA pull down assays we also tested for interactions between the EPases. Of the four EPases we studied and all possible combinations tested, the only interactions we found were between MepS-MepM and PBP4-PBP7 (Fig. S2c and S3b).

### NlpI scaffolds trimeric complexes between different EPases

Since NlpI bound multiple EPases, we tested whether NlpI could also form trimeric complexes with them. As a starting point, we tested if NlpI could scaffold MepS and PBP4 in a fixed concentration MST assay. In the presence of 3 μM NlpI the normalized fluorescence (FNorm) of fl-MepS increased, confirming the interaction between NlpI and MepS (Fig. 2c). In contrast, fl-MepS did not interact with PBP4, even when that was used in excess (30 μM; Fig. 2c). Interestingly, fl-MepS was able to bind to a saturated NlpI-PBP4 complex indicating the formation of a trimeric complex between NlpI, PBP4 and MepS (Fig. 2c). NlpI pre-incubated with excess BSA did not give the same increase in fl-MepS signal, indicating that the FNorm increase was specific to the binding of NlpI-PBP4 (Fig. 2c). We also tested whether NlpI was able to scaffold MepS and PBP7. Fl-MepS could bind pre-incubated NlpI-PBP7 complexes indicating that NlpI can scaffold both EPases and likely has different binding sites for MepS and PBP7 (Fig. 2d). Using a three component Ni^2+^-NTA pull down assay we were also able to resolve a NlpI-mediated complex containing PBP7 and MepS (Fig. S3c). The trimeric complexes were not due to direct interactions between the EPases (Fig. S2c and S3b), but rather due to NlpI scaffolding both EPases simultaneously. Thus, NlpI can scaffold at least two different trimeric EPase complexes, with MepS-PBP4 and MepS-PBP7.

### NlpI affects endopeptidase activity of MepS and MepM *in vitro*

Although NlpI interacted with and complexed several EPases, the cellular role of such complexes remained unclear. Hence, we investigated whether NlpI stimulated or repressed the activity of these EPases using *in vitro* PG digestion assays with purified sacculi or pre-digested muropeptides. EPases cleave the peptide bond between neighbouring peptides, resulting in a decrease in TetraTetra (bis-disaccharide tetrapeptide) muropeptides. Therefore, we quantified the remaining cross-linked PG substrate following incubation with the respective EPase and used the decrease in TetraTetra as an indication of EPase activity (Fig. 3a and S4a). Our results show that NlpI reduced the activity of MepM, which was more active on its own against sacculi. In contrast, MepS was inactive against sacculi and pre-digested muropeptides and that the addition of NlpI had negligble effect on MepS activity against muropeptides (Fig 3a). We did not observe significant differences in the activity of PBP4 or PBP7 in the presence or absence of NlpI (Fig. 3a). These results suggest that NlpI is able to modulate the activity *in vitro* of certain (e.g., MepM), but not all, EPases.

**Figure 3.**
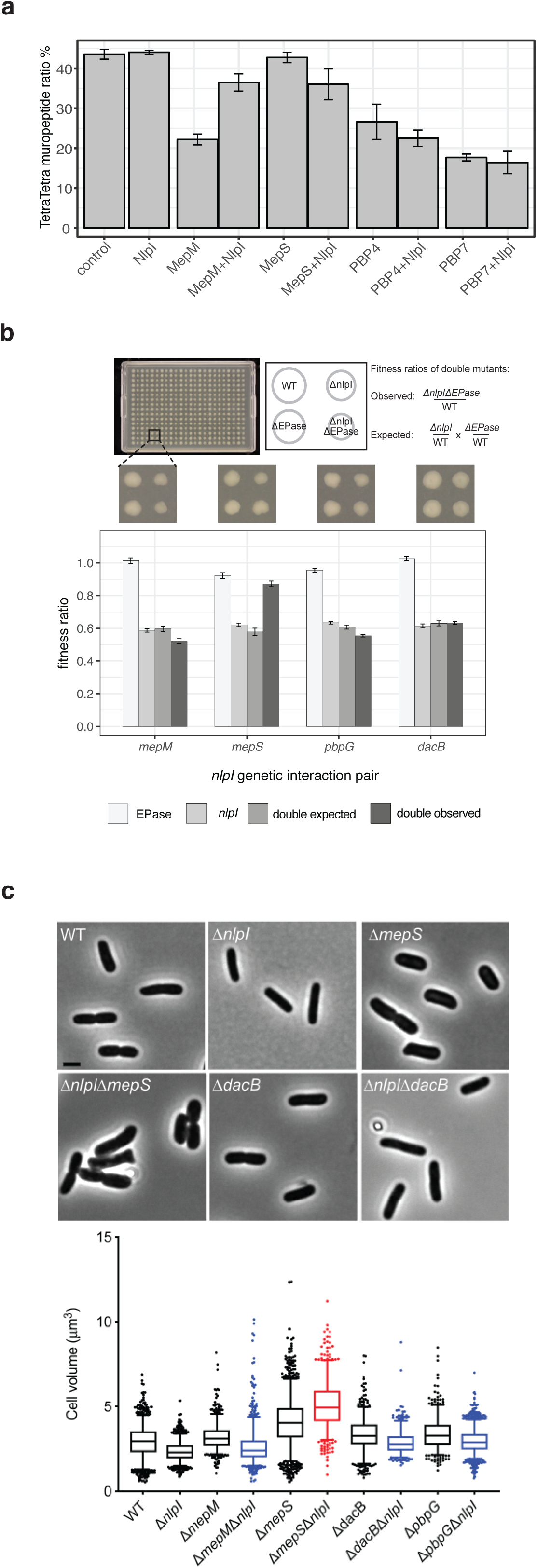
NlpI genetically interacts with MepS and affects the enzyme activity of MepS and MepM. **a.** HPLC-based PG digestion assay representing EPase activity. The graph shows the relative percentage of TetraTetra muropeptides present at the end of the respective incubation periods for each protein as described in materials and methods Values are mean ± standard deviation of three independent experiments. Representative chromatograms are shown in Fig. S4. **b.** Genetic interactions of *nlpI* with EPases. Strains were arrayed using a Rotor HDA replicator on Lennox LB agar plates and incubated for 12 h at 37°C. Each plate contained 384 colonies, 96 from the wildtype, single mutants and double mutants. Double mutants were made twice, swapping the resistance markers to the two single mutants. Colony size was quantified as a fitness readout, using the image analysis software Iris (Kritikos *et al.*, 2017). Bar plots shows the averaged values of 2 experiments (i=2, n=192). Full results can be found at supplementary table 5. **c.** *nlpI* deletion exacerbates or ameliorates the morphology of EPase-mutant strains. The graph shows the cell volume of single and double deletions strains (800 < n < 2000 cells). Shown in red are combinations where Δ*nlpI* enhances, and in blue combinations where Δ*nlpI* reduces the effect of the single EPase mutation on the morphology of the cells. Below the graph are representative images of cells lacking MepS or PBP4 in combination with a deletion of NlpI. The scale bar equals 2 µm. Gene encoding protein legend: *nlpI* encodes NlpI, *dacB* encodes PBP4, *pbpG* encode PBP7, *mepM* encodes MepM, *meps* encodes MepS.

### NlpI genetically interacts with EPases and its absence alters cell morphology

To address whether NlpI-EPase complexes are relevant for fitness in *E. coli*, we deleted *nlpI* in combination with different EPases and compared the fitness of the double mutants with that of the parental single mutants (Fig. 3b). Only *nlpI* and *mepS* exhibited a strong positive genetic interaction with the double mutant Δ*nlpI*Δ*mepS* growing as well as the Δ*mepS* mutant, and better than the Δ*nlpI* mutant. The other mutant pairs exhibited no to very mild genetic interactions (Fig. 3b). To investigate whether any of these genetic interactions were also manifested in morphological changes, all NlpI-EPase single and double mutants were grown exponentially and their cell volume was assessed using phase contrast microscopy (Fig. 3c). The Δ*nlpI* cells were 12% thinner than the wildtype cells and had a reduced cell volume. This feature was dominant in all Δ*nlpI*ΔEPase double mutants tested, except for Δ*nlpI*Δ*mepS*. The Δ*mepS* single mutant was 17% thicker compared to WT cells, resulting in an increased cell volume. This effect was enhanced in the Δ*nlpI*Δ*mepS* mutant, which resulted in 7% longer and 30% wider cells, with the highest cell volume observed of all strains which were tested (Fig. 3c). This result is intriguing, as it implies that the decreased fitness of Δ*nlpI* is associated with the presence of more MepS in the cells, which decreases their cell width. The Δ*nlpI*Δ*mepS* mutant resolves both these problems, and thus has increased fitness. Yet it still has pronounced morphological defects and retains many of the global changes in protein abundance and stability of the Δ*nlpI* mutant (Fig. S1). Overall, this indicates that the role of NlpI goes beyond the previously described proteolytic regulation of MepS (Singh *et al.*, 2015), and is probably not limited to EPases only.

### NlpI localizes along the entire cell envelope

To understand the physiological role of NlpI beyond its interaction with EPases, we investigated its cellular localization using antibodies specific for NlpI. NlpI localized in the entire envelope and not specifically at mid cell (Fig. 4a and Fig. S5), in contrast to what its interaction with divisome proteins suggested (Fig. 1d). In addition, its concentration remained constant during the cell cycle (Fig. S5a). To control for epitope occlusion by interaction partners of NlpI, we localized a functional C-terminal fusion of NlpI with a HA-tag expressed from a plasmid in the Δ*nlp*I strain. The NlpI-HA localization pattern was identical to that of NlpI (Fig. S5d). We noticed that the localization pattern of NlpI was reminiscent of the PBP synthases PBP1A and PBP2 (Banzhaf *et al.*, 2012; den Blaauwen *et al.*, 2003). Together with the links of NlpI to PG synthases observed in TPP and pull-downs (Fig.1), this made us wonder whether NlpI-EPase complexes can be part of PG machineries.

**Figure 4.**
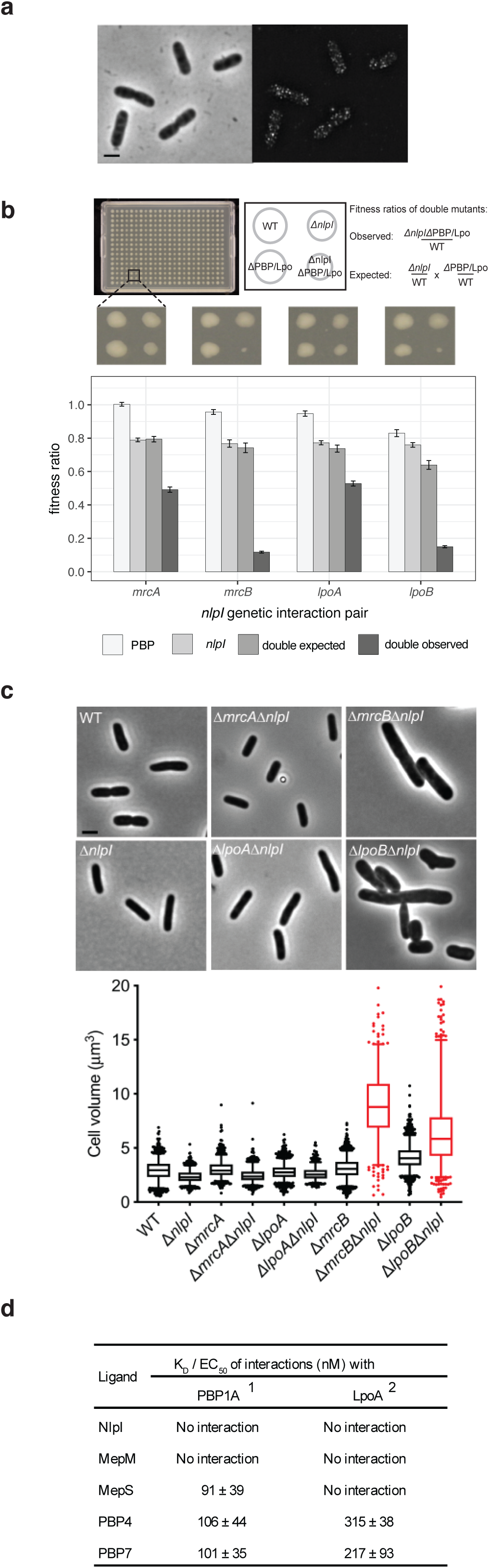
NlpI localizes along the entire cell envelope and associates with PG machineries. **a.** Phase contrast image and corresponding fluorescence SIM image of BW25113 cells that have been grown in LB at 37°C and immunolabelled with specific antibodies against NlpI. Scale bar equals 2 µm. Please see Figure S5 for further details. **b.** Genetic interactions of NlpI with PG machineries. Strains were arrayed and assessed as in Fig. 3b. Bar plots shows the averaged values of 2 experiments (i=2, n=192). Full results can be found at supplementary table 5. **c.** NlpI deletion exacerbates the morphological defects of the PBP1B/LpoB-mutant strains. The graph shows the volume of single and double deletions strains (800 < n < 2000 cells). In red are those combinations where Δ*nlpI* enhances the cell volume. The graph below shows representative images of the strains. The scale bar equals 2 µm. Gene encoding protein legend: *nlpI* encodes NlpI, *mrcA* encodes PBP1A, *lpoA* encodes LpoA, *mrcB* encodes PBP1B, *lpoB* encodes LpoB. **d.** Dissociation constants for interactions between PBP1A and LpoA with NlpI, MepM, MepS, PBP4 and PBP7 as determined by MST. The values are mean ± standard deviation of three independent experiments. ^1^PBP1A was used as fluorescently labelled protein in all assays. ^2^LpoA was used as unlabelled ligand in all combinations, except with MepM and PBP4. Binding curves are shown in Fig. S6.

### NlpI associates with PG machineries

When probing for genetic interactions, we deleted *nlpI* in combination with different PBPs and their regulators (Lpos) and compared the fitness of the double mutants with that of the parental single mutants (Fig. 4b). We noticed an almost synthetic lethality with Δ*mrcB* (encodes PBP1B) and Δ*lpoB* (encodes LpoB); Δfitness ratio of −0.62 and −0.49, respectively. Δ*mrcA* (encodes PBP1A) and Δ*lpoA* (encodes LpoA) exhibited also strong negative interactions with Δ*nlpI*; Δfitness ratio of −0.30 and −0.21 respectively (Fig. 4b). To analyse whether these strong negative genetic interactions were also reflected in the morphology of the cells, all single and double mutants were grown exponentially and imaged by phase contract microscopy. Combining Δ*nlpI* with Δ*mrcB* or Δ*lpoB* led to abnormal cell morphologies, with cells being 30% wider and up to 80% longer (Fig. 4c). This suggests that the NlpI-EPase complexes might be important for facilitating the formation of the PBP1A-mediated PG machinery. This would be consistent with the changes in thermostability of PBP1A and LpoA in Δ*nlpI* cells (Fig. 1b). Thus, we next tested the *in vitro* interactions between NlpI and respective EPases with PBP1A and LpoA. We discovered that PBP1A did not directly interact with NlpI but interacts with low nanomolar range affinities with different EPases, including MepS (apparent K_D_ = 91 ± 39 nM), PBP4 (106 ± 44 nM) and PBP7 (101 ± 35 nM) (Fig. 4d and S6a). PBP4 (315 ± 38 nM) and PBP7 (217 ± 93 nM) bound also to LpoA at slightly higher nanomolar ranges (Fig. 4d and S6b). These interactions between PG synthases and EPases would allow for PG multienzyme complexes to exist as postulated by Hӧltje (Hӧltje, 1998).

### NlpI is part of a PG multi-enzyme complex with PBP4 and PBP1A/LpoA

To further understand the interaction between PG hydrolases and synthases, we characterized in detail the interactions between PBP4 with PBP1A/LpoA and NlpI by MST. We used a fixed concentration MST assay to show that fluorescently labelled PBP1A and LpoA are able to bind a preformed PBP4-NlpI complex (Fig. 5a and 5b). Whilst the binding of PBP4 and PBP4-NlpI to fl-PBP1A resulted in an increase in FNorm values (which was not the case in the presence of NlpI alone); binding of PBP4 and PBP4-NlpI to LpoA consistently resulted in an enhanced initial fluorescence. This indicated that the ligand was binding in close proximity to the probe and was affecting the local environment of the fluorophore and subsequently its fluorescence yield. Since the change in fluorescence was a property of ligand binding (Fig. S8b), the raw fluorescence data as opposed to the FNorm values were plotted in this instance (Fig. 5b). These consistent increases in fluorescence reflect the binding of PBP4 and PBP4-NlpI to LpoA and suggest that the presence of NlpI does not prevent the interaction of PBP4 with LpoA (Fig. 5b). To our knowledge, this is the first evidence that PG-synthases and PG-hydrolases form multienzyme complexes with regulatory lipoproteins to possibly coordinate PG-synthesis in Gram-negative bacteria.

**Figure 5.**
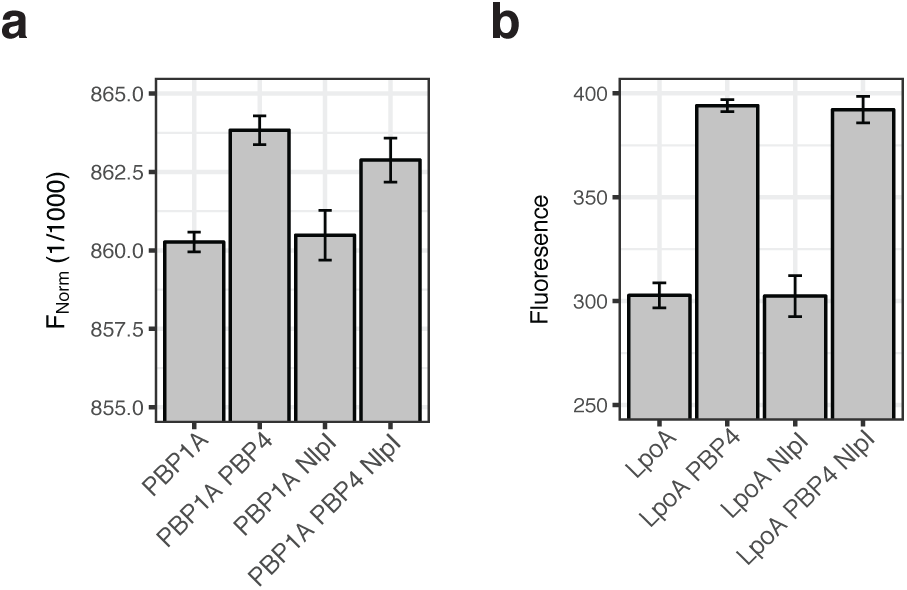
PBP4 forms a PG multienzyme complex with NlpI and PBP1A/LpoA. **a-b.** PBP4 has different interaction sites for PBP1A/LpoA and NlpI as shown by a single concentration MST assay. Unlabelled PBP4 and excess NlpI were titrated against fluorescently labelled PBP1A and LpoA and changes in FNorm or fluorescence are plotted. Values are mean ± standard deviation of three independent experiments.

## Discussion

*E. coli* contains a repertoire of more than 20 periplasmic hydrolases providing specificity to almost every bond present in PG (Chodisetti & Reddy, 2019; Singh *et al.*, 2012; van Heijenoort, 2011; Vollmer *et al.*, 2008b; Yunck *et al.*, 2016). However, with the exception of amidases, it is unclear how these hydrolases are regulated to prevent autolysis (Uehara et al., 2009). This study identifies NlpI as a novel scaffolding protein of EPases that might co-ordinate hydrolases within PG synthesis machineries. NlpI seems to also bind to several other hydrolytic enzymes, including some members of the amidase and LTase families. The details of these interactions will be investigated in future work.

### Deletion of *nlpI* impacts envelope biogenesis beyond the proteolytic regulation of MepS levels

NlpI interacts with MepS and targets it for digestion via the protease Prc (Singh *et al.*, 2015). Δ*nlpI* increases the abundance of MepS (Fig. 1a) along with conferring an OM hypervesiculation phenotype (Schwechheimer *et al.*, 2015; Singh *et al.*, 2015). The hypervesiculation is suppressed in Δ*nlpI*Δ*mepS* mutants, indicating that it is caused by the increased MepS levels (Schwechheimer *et al.*, 2015). Similarly, we show that unregulated MepS is also part of the fitness defect of Δ*nlpI*, as the double Δ*nlpI*Δ*mepS* mutant has most of its fitness defect restored. Yet, there are several pieces of evidence that NlpI has additional functions in addition to the proteolytic degradation of MepS.

First, the cellular morphology of the Δ*nlpI*Δ*mepS* mutant is not restored. The Δ*nlpI* and the Δ*mepS* mutant are thinner and thicker, respectively, compared to wildtype cells; which is in line with MepS playing an important role for cell elongation (Singh *et al.*, 2012) and Δ*nlpI* cells having this activity uncontrolled. However, the morphology of the double mutant is exacerbated compared to an Δ*mepS* mutant, with the cells becoming not only thicker but also more elongated (Fig. 3c). This strongly implies that NlpI has additional functions to the proteolytic degradation of MepS. Second, our biochemical evidence (TPP, affinity chromatography) suggest that NlpI binds to and affects a number of PG related processes, including both PG hydrolysis and synthesis enzymes and their regulators. NlpI seems to bind strongly to amidases and their regulators (AmiA, EnvC; Fig. 1b, d), LTases (MltA, MltC; Fig. 1b,d) and EPases (MepM, MepS, PBP4, PBP7; Fig 2, S2, S3) in the context of PG biosynthetic machineries (Fig. 4-6). This raises the possibility that NlpI scaffolds, or even regulates, several classes of hydrolases beyond its function towards EPases. Moreover, to the best of our knowledge, this is the first evidence that NlpI has additional functions during PG synthesis.

### NlpI interacts with several EPases at physiologically relevant concentrations

Immobilized NlpI retained the EPases PBP4 and PBP7, raising the possibility that NlpI interacts with additional EPases along with MepS (Fig. 1c). Especially since MepS was not part of the proteins being pulled down, and is known to bind to NlpI, we decided to investigate this further. Using MST and pull-down assays, we validated interactions between NlpI and 3 other DD-EPases; MepM, PBP4 and PBP7, all of them with apparent K_D_ or EC_50_ values in the nanomolar range (Fig. 2c and S2a). We estimated the concentration of these proteins in the periplasm, assuming cell dimensions of 4.77 × 10^−6^ m (length) and 1.084 × 10^−6^ m (diameter), with a periplasmic width of 21 × 10^−9^ m (Banzhaf *et al.*, 2012; Beveridge, 1995) (Fig. 6a). We conclude that the NlpI-EPases interactions identified in the present work are all, in principle, able to occur in the cell (Fig. 6a). Furthermore, our data showed that NlpI could also affect the activity of some of these EPases; for example, the activity of MepM against intact sacculi, was reduced in the presence of NlpI (Fig. 3a). Further, as NlpI facilitates to proteolytic degradation of MepS (Singh *et al.*, 2015), NlpI could be generally restricting the role of elongation EPases (Singh *et al.*, 2012).

**Figure 6.**
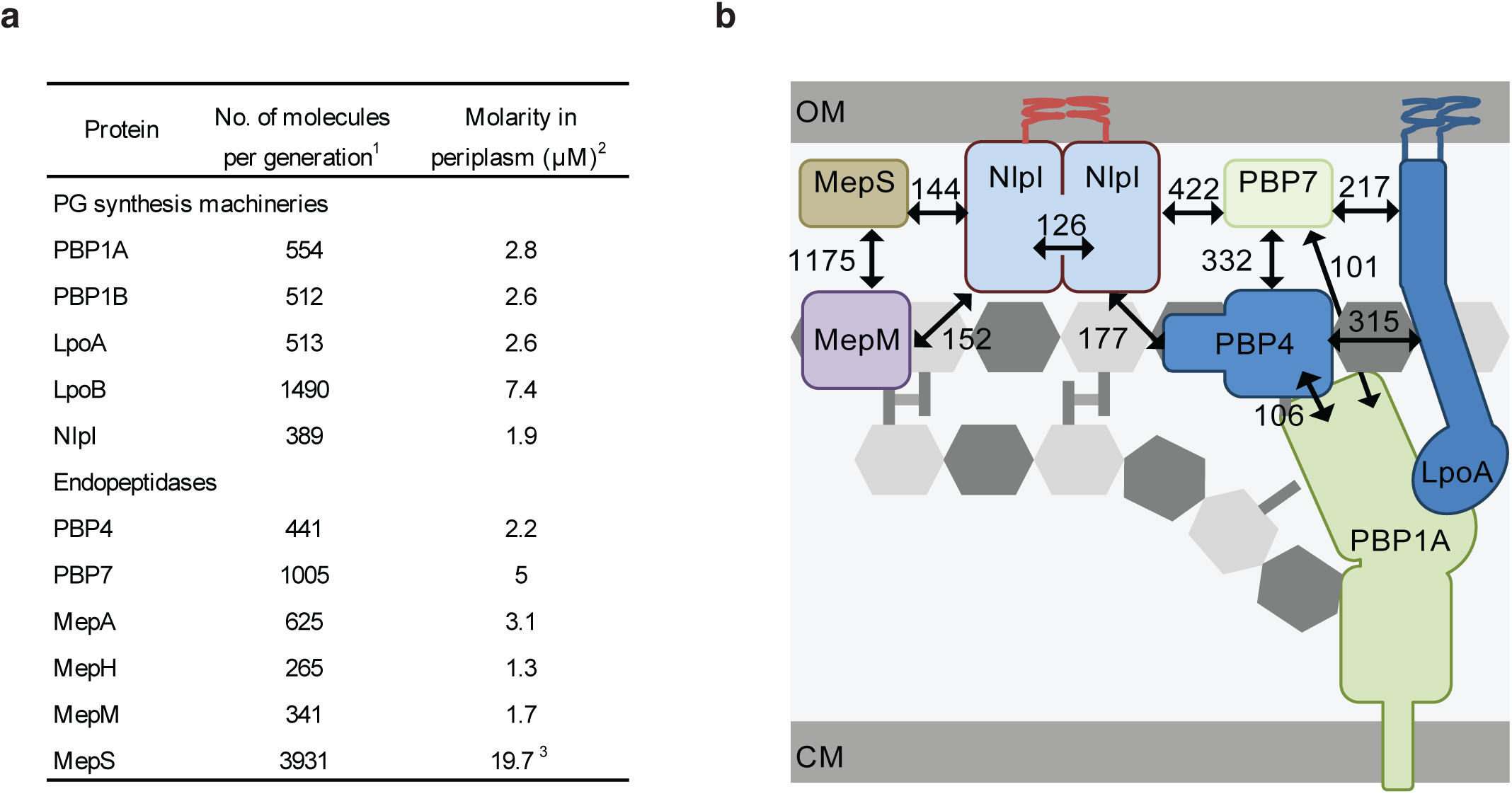
Proposed model for a role of NlpI in nucleating formation of PG multi-enzyme complexes containing EPases. **a.** Estimated number of molecules and molarity of PG synthesis machineries/regulators and EPases. ^1^Numbers obtained by ribosomal profiling in rich growth medium (Li *et al.*, 2014). ^2^Concentration of monomer. ^3^Decreases in the presence of NlpI (Singh *et al.*, 2015). The periplasmic concentrations of proteins were estimated for a cell with periplasmic volume of 3.33 ×10^−16^ L, where 1 molecule corresponds to 5 nM. **b.** Hypothetical model of NlpI scaffolding endopeptidases during cell elongation. Black arrows indicate interacting proteins with numbers indicating apparent EC_50_ / K_D_ values. OM, outer membrane; CM, cytoplasmic membrane. MepS-PBP1A interaction is not represented due to illustrative restrictions.

With regards to activity of EPases, we note that we were unable to observe the DD-EPase activity of MepS, previously reported in (Singh *et al.*, 2012) (Fig. 3a). However, while addition of NlpI had no effect on the activity of PBP4 or PBP7, there was a very slight stimulation of MepS activity against isolated muropeptides in the presence of NlpI, following overnight incubation (Fig. 3a). Overall, these results raise the possibility that NlpI could modulate the activity of specific hydrolases along with its role as a scaffolding protein. Since NlpI seems able to bind several of the hydrolytic enzymes simultaneously (at least different EPases); its role in the regulation of activity may become clearer when probed in the context of these multimeric complexes.

### NlpI scaffolds multiprotein complexes with PG hydrolytic enzymes within the context of PG biosynthesis machineries

NlpI is able to form trimeric complexes with different EPases that lack mutual interactions. Examples of such complexes resolved in the present work are MepS-NlpI-PBP4 and MepS-NlpI-PBP7 (Fig. 2c and 2d). Since NlpI has four helix-turn-helix TPR-like repeats per monomer, it remains to be seen whether the different TPR helixes are specific for different binding partners (Wilson *et al.*, 2005) and/or different type of hydrolytic enzymes. Nevertheless, the ability of NlpI to bind multiple ligands simultaneously would be consistent with the idea that TPR domains facilitate the formation of multi-protein complexes (Blatch & Lassle, 1999; Cortajarena & Regan, 2006). In this sense, NlpI is more promiscuous in nature than the previously identified amidase regulators EnvC and NlpD, which appear to have specificity to their cognate amidases (Uehara *et al.*, 2009). Despite binary interactions between various EPases and PBP1A/LpoA being able to occur in the absence of NlpI (Fig. 4d), we hypothesize that NlpI could also sequester additional or specific sets of EPases and other hydrolytic enzymes, determining the specificity of such synthetic machineries. Accordingly, our finding that PBP4 is able to simultaneously bind PBP1A/LpoA and NlpI supports the idea that NlpI could specifically scaffold hydrolases at active PG synthases (Fig. 5a and 5b). The ability of an OM-anchored NlpI to complex EPases and other hydrolases would not only serve to locally concentrate those enzymes near PG-synthesis complexes, but also to maintain the active hydrolases in the space between the PG layer and OM; facilitating cleavage of the mature PG of the sacculus and keeping them at distance to the newly synthesized PG, which emerges between the CM and PG layer and is not subject to turnover. NlpI molecules are outnumbered by the amount of potential binding partners in the periplasm, so it is unlikely that there is an abundance of free NlpI (Fig. 6a). EPase regulation might occur on the level of binding affinity to NlpI and its TPR-like domains. This would see NlpI resembling a “dock” for EPases (and possibly other hydrolases) to make them available for PG synthesis complexes when needed. Such a system would allow for greater flexibility, as NlpI interacts with many hydrolases. In contrast, the specificity could be encoded on the level of the hydrolases. As demonstrated, EPases interact directly with PG-synthases, but those interactions might be specific to them particular EPases (and no other hydrolases) and/or might be subject to environmental cues or to competition for the same binding site. Therefore NlpI could be a more general adaptor of hydrolases, as suggested by its interactions with amidases and LTases (Fig. 1b, d), bringing a set of hydrolases to biosynthetic complexes. This hypothesis will require more work in the future to ascertain.

Around 20 years ago, Hӧltje hypothesized that growth of the PG sacculus requires both synthases and hydrolases working in tandem to enable a safe and coordinated enlargement (Hӧltje, 1998). However, it has also been suggested that EPases are not necessarily part of multi-protein complexes; as overproduction of three different EPases confers mecillinam resistance (Lai *et al.*, 2017). In this work, we provide the first evidence of interactions between PBP1A/LpoA with PBP4 and hypothesize that interactions between NlpI and other EPases could facilitate their delivery to PG-synthesis complexes during PG growth. The existence of PG multi-protein complexes is not necessarily contrasting the idea that EPases and/or NlpI-EPase complexes may in part localize outside of such PG assembly machineries. This work, and the work of others supports the idea that PG multi-protein complexes are highly dynamic and driven by transient protein-protein interactions (Pazos *et al.*, 2017). In addition, the existence of such PG multi-protein complexes is in line with the recent isolation of an 1 MDa cell division complex (Trip & Scheffers, 2015).

### NlpI functions together with the PBP1A/LpoA PG machinery

We studied the localization of NlpI to infer if NlpI scaffolds complexes exclusively for cell elongation or division. The localization pattern of NlpI is spotty and diffusive with no enrichment at the mid-cell (Fig 4a). NlpI was previously shown to be located in the OM of bacterial cells, containing an N-terminal cysteine residue at position 19 that is likely the target for lipoprotein modification (Ohara *et al.*, 1999; Teng *et al.*, 2010). The subsequent N-acyl-S-sn-1,2-diacylglyceryl-cysteine is processed, culminating in the tethering of the mature protein to the inner leaflet of the OM (Noland et al., 2017; Wilson et al., 2005). It is hence also possible that interactions between NlpI and hydrolases concentrate and facilitate cleavage from the outer face of the PG layer. Its disperse localization would enable binding of EPases involved in both division and elongation. NlpI was shown to bind a number of essential divisome proteins at high salt concentrations in our affinity chromatography experiment (Fig 1d), suggesting that it is at least a transient member of the divisome. In addition, the negative genetic interaction of *nlpI* with *mrcA* (PBP1A) and *mrcB* (PBP1B) raises the possibility that NlpI can affect both the elongasome and the divisome. This is in line with both PBP1B/LpoB and PBP1A/LpoA complexes showing changes in thermostability in Δ*nlpI* cells (Fig. 1b). However, because the genetic interactions of *nlpI* with *mrcB* (PBP1B) were stronger and led to a near synthetic lethality, we reasoned that NlpI predominantly worked with the PBP1A/LpoA machinery. Cells lacking PBP1B depend on a functional PBP1A/LpoA complex to achieve growth (Yousif *et al.*, 1985). An alternative scenario is that cells with only PBP1A/LpoA are more sensitive to genetic and chemical perturbations in the cell wall than cells with only PBP1B/LpoB, because latter is more efficient. Although we found no direct interaction between NlpI with PBP1A or LpoA by MST assay (Fig. 4d and S6), a complex of NlpI-PBP4-PBP1A could be formed with PBP4 as the linking protein (Fig. 5). The multitude of interactions between PBP1A/LpoA, different EPases and NlpI (Fig. 4d and S6) could enable the formation of different active synthase-hydrolase complexes under a range of growth conditions or availability of particular proteins (Fig. S6) (Pazos *et al.*, 2017).

In conclusion, this work provides the first evidence for NlpI as a novel adaptor of EPases, and possibly other classes of PG hydrolases. NlpI is likely involved in co-ordinating PG-multienzyme complexes by way of nucleating complexes between synthases, EPases and NlpI (Fig. 6b).

## Materials and methods

### Media and Growth conditions

Strains used in this work were grown in LB media (1% tryptone, 1% NaCl, 0.5% yeast extract) at 37°C, unless otherwise stated. Antibiotics were used at the following concentrations (µg/ml): ampicillin (Amp), 100; chloramphenicol (Cam), 25; kanamycin (Kan), 50. MC4100 cells were grown to steady state (Vischer et al., 2015) in glucose minimal medium containing 6.33 g of K_2_HPO_4_.3H_2_O, 2.95 g of KH_2_PO_4_, 1.05 g of (NH_4_)_2_, 0.10 g of MgSO_4_.7H_2_O, 0.28 mg of FeSO_4_.7H_2_O, 7.1 mg of Ca(NO_3_)_2_.4H_2_O, 4 mg of thiamine and 4 g of glucose. For strain MC4100, 50 µg lysine per L was added. Absorbance was measured at 450 nm with a 300-T-1 spectrophotometer (Gilford Instrument Laboratories Inc.).

### Bacterial Strain Construction

BW25113 was used as the parent strain (WT) for this study. Strains were generated, unless otherwise stated, by transducing P1 lysates derived from the corresponding deletion strains of the Keio and Aska strain collections (Adams & Luria, 1958; Baba *et al.*, 2006).

#### Generation of the the NlpI-HA tagged strain

For HA-tagging, pKD13 (kanamycin resistant) was used as a PCR template. The kanamycin cassette was amplified by PCR with the primers 74-NlpI-HA-O1 and 87-NlpI-HA-O2. The primer 74-NlpI-HA-O1 was carrying from 5’ to 3’: the homology region of the C-terminal of NlpI (without the STOP codon), 2 HA-tag and the homology region of the N-terminus of the kanamycin cassette (from the pKD13). The primer 87-NlpI-HA-O2 was carrying from 5’ to 3’ the homology region of the downstream region of NlpI and the homology region of the C-terminus of the kanamycin cassette.

#### Generation of the NlpI-Strep-Flag (-SF) tagged strain

For SF-tagging, pJSP1 (containing the SF-tag and a kanamycin cassette) was used as a PCR template. The SF-tag and kanamycin cassette were amplified by PCR with the primers 175-NlpI-SF-O1 and 176-NlpI-SF-O2. The primer 175-NlpI-SF-O1 was carrying from 5’ to 3’ the homology region of the C-terminal of NlpI (without the stop codon) and the homology region of the N-terminal of the Strep-Flag tag (from the pJSP1). The primer 176-NlpI-SF-O2 was carrying from 5’ to 3’ the homology region of the downstream region of NlpI and the homology region of the C-terminus of the kanamycin cassette.

BW25113 transformants carrying a Red helper plasmid were grown in 5-ml SOB cultures with ampicillin and l-arabinose at 30°C to an OD_600_ of ≈0.6 and then made electrocompetent by concentrating 100-fold and washing three times with ice-cold 10% glycerol. PCR products were gel-purified, digested with *Dpn*I, repurified, and suspended in elution buffer (10 mM Tris, pH 8.0). Electroporation was done by using a Cell-Porator with a voltage booster and 0.15-cm chambers according to the manufacturer’s instructions (GIBCO/BRL) by using 25 μl of cells and 10–100 ng of PCR product. Shocked cells were added to 1 ml SOC, incubated 1 h at 37°C, and then one-half of the incubation/cells were spread onto agar to select Km^R^ transformants. They were colony-purified once non-selectively at 37°C and then tested for ampicillin sensitivity to test for loss of the helper plasmid.

### Eliminating Antibiotic Resistance Gene for the NlpI-HA

pCP20 is an ampicillin and Cm^R^ plasmid that shows temperature-sensitive replication and thermal induction of FLP synthesis (Cherepanov & Wackernagel, 1995). Km^R^ mutants were transformed with pCP20, and ampicillin-resistant transformants were selected at 30°C, after which a few were colony-purified once non-selectively at 43°C and then tested for loss of all antibiotic resistances. The majority of the mutants lost the FRT-flanked resistance gene and the FLP helper plasmid simultaneously.

### Immunolabeling

After reaching steady state, the cells were fixed for 15 min by addition of a mixture of formaldehyde (f.c. 2.8%) and glutaraldehyde (f.c. 0.04%) to the shaking water bath and immunolabeled as described (Buddelmeijer et al., 2013) with rabbit polyclonal antibodies against NlpI or against the HA-tag. As secondary antibody, donkey anti-rabbit conjugated to Cy3 or conjugated to Alexa488 (Jackson Immunochemistry, USA) diluted 1:300 in blocking buffer (0.5% (wt/vol) blocking reagents (Boehringer, Mannheim, Germany) in PBS) was used, and the samples were incubated for 30 min at 37°C. For immunolocalization, cells were immobilized on 1% agarose in water slabs coated object glasses as described (Koppelman et al.,2004) and photographed with an Orca Flash 4.0 (Hamamatsu) CCD camera mounted on an Olympus BX-60 fluorescence microscope through a 100x/N.A. 1.35 oil objective. Images were taken using the program ImageJ with MicroManager (https://www.micro-manager.org). SIM images were obtained with a Nikon Ti Eclipse microscope and captured using a Hamamatsu Orca-Flash 4.0 LT camera. Phase contrast images were acquired with a Plan APO 100x/1.45 Ph3 oil objective. SIM images were obtained with a SR APO TIRF 100x/1.49 oil objective, using 3D-SIM illumination with a 488 nm laser, and were reconstructed with Nikon-SIM software using the values 0.23-0.75-0.10 for the parameters Illumination Modulation Contrast (IMC), High Resolution Noise suppression (HNS) and Out of focus Blur Suppression (OBS).

### Image analysis

Phase contrast and fluorescence images were combined into hyperstacks using ImageJ (http://imagej.nih.gov/ij/) and these were linked to the project file of Coli-Inspector running in combination with the plugin ObjectJ (https://sils.fnwi.uva.nl/bcb/objectj/). The images were scaled to 15.28 pixel per μm. The fluorescence background has been subtracted using the modal values from the fluorescence images before analysis. Slight misalignment of fluorescence with respect to the cell contours as found in phase contrast was corrected using Fast-Fourier techniques as described in (Vischer *et al.*, 2015). Data analysis was performed as described in (Vischer *et al.*, 2015). In brief, midcell was defined as the central part of the cell comprising 0.8 µm of the axis. From either cell part, midcell and remaining cell, the volume, the integrated fluorescence, and thus the concentration of fluorophores was calculated. The difference of the two concentrations is multiplied with the volume of midcell. It yields FCPlus (surplus of fluorescence). For age calculation, all cell lengths are sorted in ascending order. Then the equation:

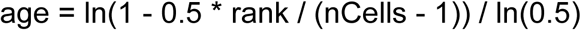

is used, where *rank* is a cell’s index in the sorted array, *nCells* is the total amount of cells and *age* is the cell’s age expressed in the range 0 to 1.

### Ni^2+^-NTA pulldown assay

His-tagged proteins of interest were incubated with untagged or native ligands, in the presence of Ni^2+^-NTA coupled agarose beads (Qiagen). Beads were pre-equilibrated with dH_2_O and binding buffer (10 mM HEPES/NaOH, 10 mM MgCl_2_, 150 mM, NaCl 0.05% Triton X-100, pH 7.5) by centrifugation at 4000 × *g*, 4 min at 4°C. Samples were incubated overnight on a spinning plate at 4°C before beads were washed 3-6 times with 10 mM HEPES/NaOH, 10 mM MgCl_2_, 150 mM, NaCl 0.05% Triton X-100, 30 mM Imidazole, pH 7.5. Retained material was eluted from Ni^2+^-NTA beads using proteus spin columns and boiling at 100°C in SDS-buffer (50 mM Tris/HCl pH 6.8, 2% SDS, 10% glycerol 0.02% bromophenol blue, 10% β-mercaptoethanol). Elutions were diluted 1:1 with dH_2_O and proteins were separated by SDS-PAGE for analysis.

### Protein overexpression and purification

Prior to purification, plasmids of interest were transformed into *E. coli* strain BL21 (λDE3) and grown overnight in LB agar (1.5% w/v) containing appropriate antibiotic, at 37°C. Transformants were inoculated into 50 ml of LB with appropriate antibiotic and grown overnight at 37°C, shaking. Pre-cultures were diluted 1:40 in 2 L LB and grown to OD_578_ 0.5-0.6, at 37°C. Induction conditions are specified for each respective protein below. After overexpression, cells were harvested by centrifugation at 7500 × *g*, 15 min, 4°C. Pellets were re-suspended in buffer I (25 mM Tris/HCl, 300 mM NaCl, pH 7.5) with the addition of a small amount of DNase (Sigma) and 100 μM P.I.C and PMSF. Cells were lysed by sonication (Branson digital) and the lysate was centrifuged at 14000 × *g*, 1 h, 4°C, before the supernatant was applied at 1 ml/min to a 5 ml chromatography column attached to an ÄKTA Prime plus (GE Healthcare).

If desired, the removal of his-tags for tagged constructs, following IMAC steps, was achieved by incubating protein samples with 1 unit/ml of restriction grade thrombin (Novagen). This was carried out overnight at 4°C in 25 mM Tris/HCl, 200 mM NaCl, pH 8.0 or 25 mM HEPES/NaOH, 300 mM NaCl, 10% glycerol, pH 7.5, depending on the next purification step. Removal of His-tag was verified by western blot with monoclonal α-His –HRP (1:10000) antibody (Sigma).

### Purification of MepM

MepM was purified as previously described in (More *et al.*, 2019).

### Purification of MepS and MepS(C68A)

MepS and MepS(C68A) overexpression was induced with 1 mM IPTG for 90 min at 37 °C. Following harvesting, lysate was applied to a 5 ml HisTrap HP column (GE healthcare) in buffer containing 25 mM Tris/HCl, 300 mM NaCl, 20 mM Imidazole pH 7.5. Protein was eluted in 25 mM Tris/HCl, 300 mM NaCl, 400 mM imidazole, 10% glycerol, pH 7.5. Protein purity and yield were analyzed by SDS PAGE and the fractions of interest were pooled and dialyzed overnight against 25 mM HEPES/NaOH, 300 mM NaCl, 10% glycerol, pH 7.5. Protein was concentrated to ~ 5ml using Vivaspin concentrator spin columns (Sartorius) at 4500 × *ɡ,* 4°C and applied to a HiLoad 16/600 Superdex 200 column (GE healthcare) at 1 ml/min. Protein purity and yield were analyzed by SDS-PAGE and the best fractions were pooled and stored at −80°C.

### Purification of NlpI

NlpI overexpression was induced with 1 mM IPTG, 3 h at 30°C, before harvesting of cells as described above. Following harvesting, lysate was applied to a 5 ml HisTrap HP column (GE healthcare) and washed with buffer containing 25 mM Tris/HCl, 300 mM NaCl, 20 mM Imidazole pH 7.5. Protein was eluted in 25 mM Tris/HCl, 300 mM NaCl, 400 mM imidazole, 10% glycerol, pH 7.5. Protein purity and yield was analyzed by SDS PAGE and the fractions of interest were pooled and dialyzed overnight against 25 mM HEPES/NaOH, 200 mM NaCl, 10% glycerol, pH 7.5. Protein was concentrated to ~ 5ml using Vivaspin concentrator spin columns (Sartorius) at 4500 × *ɡ,* 4°C and applied to a HiLoad 16/600 Superdex 200 column (GE healthcare) at 1 ml/min. Protein purity and yield were analyzed by SDS-PAGE and the best fractions were pooled and stored at −80°C.

### Purification of PBP4

Purification of native PBP4 followed an adapted protocol from (Kishida *et al.*, 2006). PBP4 overexpression was induced with 1 mM IPTG for 8 h at 20°C and then harvested by centrifugation at 7500 × *ɡ,* 4°C, 15 min. Cell pellets were resuspended in buffer (50 mM Tris/HCl, 300 mM NaCl, pH 8.0) and lysed by sonication before centrifuging at 14000 × *ɡ*, 1 h, 4°C and reducing NaCl concentration by stepwise dialysis in a Spectra/Por dialysis membrane (MWCO 12-14 kDa). Cell supernatant was first dialyzed against dialysis buffer I (50 mM Tris/HCl, 200 mM NaCl, pH 8.5) for 1 h at 4°C, then against diaysis buffer II (50 mM Tris/HCl, 100 mM NaCl, pH 8.5) for 1 h at 4°C and then finally against dialysis buffer III (50 mM Tris/HCl, 30 mM NaCl, pH 8.5), O/N at 4°C. Dialyzed protein sample was then centrifuged at 7500 × *ɡ,* 4°C, 10 min and supernatant applied to a 5 ml HiTrap Q HP IEX column in 25 mM Tris/HCl, 30 mM NaCl, pH 8.5. Protein was eluted from the column with a linear gradient of buffer 2 containing 25 mM Tris/HCl, 1 M NaCl, pH 8.0, over a 100 ml volume. Fractions of interest were analyzed by SDS-PAGE and the best fractions were pooled and dialyzed O/N, at 4°C, against dialysis buffer containing 10 mM Potassium phosphate, 300 mM NaCl, pH 6.8. Protein was applied at 1 ml/min to a 5 ml ceramic hydroxyapatite column (BioRad Bioscale^TM^) in the dialysis buffer. Fractionation of proteins was achieved by using a linear gradient of buffer 2 (500 mM Potassium phosphate, 300 mM NaCl, pH 6.8) over a 50 ml gradient. Fractions of highest purity and yield were dialyzed overnight against 25 mM HEPES/NaOH, 300 mM NaCl, 10% glycerol, pH 7.5 and concentrated to ~ 5ml using Vivaspin concentrator spin columns (Sartorius). Protein sample was applied to a HiLoad 16/600 Superdex 200 column (GE healthcare) at 1 ml/min pre-equilibrated with dH_2_O and buffer I (25 mM HEPES/NaOH, 300 mM NaCl, 10% glycerol, pH 7.5). Samples were analyzed by SDS-PAGE and fractions containing purified protein was pooled and stored at −80°C.

### Purification of PBP7

PBP7 overproduction was induced with 1 mM IPTG for 3 h at 30°C before being harvested by centrifugation as described above and re-suspended in buffer I (25 mM Tris/HCl, 500 mM NaCl, 20 mM Imidazole, pH 7.5). Following sonication and subsequent centrifugation as described above, the lysate was applied to a 5 ml HisTrap HP column (GE healthcare) and washed with 4 column volumes of buffer I; before bound protein was eluted with buffer II (25 mM Tris/HCl, 300 mM NaCl, 400 mM Imidazole pH 7.5). Samples were analyzed by SDS PAGE and dialyzed overnight against 25 mM HEPES/NaOH, 300 mM NaCl, 10% glycerol, pH 7.5 before being concentrated to ~ 5ml using Vivaspin concentrator spin columns (Sartorius) at 4500 × *ɡ,* 4°C. Protein samples were applied to a HiLoad 16/600 Superdex 200 column (GE healthcare) at 1 ml/min pre-equilibrated with dH_2_O and buffer I (25 mM HEPES/NaOH, 300 mM NaCl, 10% glycerol, pH 7.5). Samples were analyzed by SDS-PAGE and the purest fractions with highest yield were pooled and stored at −80°C.

### Purification of PBP1A and LpoA

Purification of PBP1A and LpoA was as described previously in (Born *et al.*, 2006) and (Jean *et al.*, 2014), respectively.

### Microscale Thermophoresis assays

NlpI, MepS, PBP1A, PBP4 and PBP7 were labelled on amines with NT647 RED-N-hydroxysuccinimide (NHS) reactive dye, whilst LpoA was labelled on cysteines with NT647 RED-Maleimide reactive dye (Nanotemper) according to manufacturer’s instructions (Nanotemper) and as described in (Jerabek-Willemsen *et al.*, 2011). Two fold serial dilution of proteins were done in MST buffer containing 25 mM HEPES/NaOH, 150 mM NaCl, 0.05% Triton X-100, pH 7.5. Unlabelled ligands were titrated from the following starting concentrations: NlpI (50 μM), MepM (30 μM), MepS(C68A) (30 µM), PBP4 (50 μM), PBP7 (30 μM) and LpoA (30 μM). Ligands were serially diluted 16 times and assayed for interactions with labelled proteins of interest at 10 - 40% MST power in standard or premium capillaries on a Monolith NT.115. Binding curves and kinetic parameters were plotted and estimated using NT Analysis 1.5.41 and MO. Affinity Analysis (x64) software.

#### SDS Denaturation (SD) test

Prior to all MST measurements, capillary scans were carried out to check for consistent initial fluorescence counts. In assays which showed concentration dependent changes in fluorescence intensity, SDS denaturation tests were carried out to investigate whether changes were non-specific or were a property of ligand binding. 10 μl from samples containing the highest and lowest concentration of unlabelled ligand were centrifuged 10 000 × *g,* 5 min, RT and mixed 1:1 volume ratio with SD-test buffer (40 mM DTT, 4% SDS). Mixtures were boiled at 100°C for 10 min to abolish ligand binding before being spun down and subjected to another capillary scan. If fluorescence intensities were back to within the margin of error, then initial changes were due to ligand binding and binding curves were plotted using raw fluorescence values.

#### Fixed ligand concentration MST assays for trimeric complexes

For fixed ligand concentration MST assays, labelled proteins were titrated against a fixed concentration of unlabelled proteins, respectively. In fixed concentration assays with labelled MepS or LpoA, unlabelled PBP4-NlpI or PBP7-NlpI complexes were preformed by incubating NlpI (3 μM) with excess PBP4 or PBP7 (30 μM), on ice for 10 min. In fixed concentration assays with labelled PBP1A, unlabelled PBP4-NlpI complex was pre-formed by incubating PBP4 (0.5 μM) with NlpI (1 μM), on ice for 10 min. Thermophoresis or fluorescence of labelled protein in the presence of unlabelled ligands was determined using NT Analysis 1.5.41 and MO Affinity Analysis (x64) software.

### Analytical ultracentrifugation

Purified NlpI was dialysed O/N against 25 mM HEPES/NaOH, 150 mM NaCl, pH 7.5, in preparation for AUC. AUC sedimentation velocity (SV) experiments were carried out in a Beckman Coulter (Palo Alto, CA, USA) ProteomeLab XL-I analytical ultracentrifuge using absorbance at 280 nm and interference optics. All AUC runs were carried out at a rotation speed of 45,000 rpm at 20°C using an 8-hole AnTi50 rotor and double-sector aluminium-Epon centrepieces. The sample volume was 400 µl and the sample concentrations ranged between 0.3 and 1.2 mg/ml. The partial specific volumes (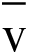) for the proteins were calculated from the amino acid sequence of NlpI, using the program SEDNTERP (Laue *et al.*, 1992). Sedimentation velocity profiles were treated using the size-distribution c(s) model implemented in the program SEDFIT14.1 (Schuck, 2000). The experimental values of the sedimentation coefficient were corrected for the viscosity and density of the solvent, relative to that of H_2_O at 20°C (s_20,w_). The atomic coordinates from the published crystal structure of (Wilson *et al.*, 2005) was used to calculate the sedimentation coefficient values for the monomer and dimer of NlpI using the program SoMo (Brookes *et al.*, 2010).

### *In vitro* PG digestion assays

PG digestion assays and subsequent muropeptide composition analysis was carried out as previously described (Glauner, 1988). 10% (v/v) substrate isolated from *E.coli* strain MC1061 was utilized in digestion reactions as follows: MepM (2 μM) ± NlpI (4 μM) incubated against intact sacculi for 4h, MepS (5 μM) ± NlpI (10 μM) incubated against muropeptides O/N, PBP4 (2 μM) ± NlpI (4 μM) incubated against sacculi for 4 h, PBP7 (2 μM) ± NlpI (4 μM) incubated against muropeptides for 4 h. Standard reaction conditions were 10 mM HEPES/NaOH, 10 mM MgCl_2_, 150 mM NaCl, 0.05% Triton X-100, pH 7.5, in 100 μl reaction volume. Following incubation, samples were boiled at 100°C, 10 min, to terminate reactions before digesting remaining PG overnight at 37°C, with 1 μM cellosyl (Hoechst, Frankfurt am Main, Germany). The samples were centrifuged at 10 000 × *g* for 5 min, RT, to obtain digested muropeptide products in the supernatant. Following digestion, muropeptide products were reduced with NaBH_4_, adjusted to pH 4-5 and separated for analysis by reversed-phase HPLC (Glauner, 1988).

Pre-digested muropeptides were obtained by incubating intact sacculi from *E. coli* strain MC1061, with 1 μM cellosyl at 37°C, overnight. Following this, reactions were terminated by boiling at 100°C for 5 min and the muropeptide substrates obtained by centrifugation at 10 000 × *g* for 5 min, RT and taking the supernatant. Reactions were then carried out and prepared for analysis as described above.

### Purification of anti-NlpI

This protocol was adapted from a previously published method (Banzhaf *et al.*, 2012). Serum against NlpI was obtained from rabbits at Eurogentec (Herstal, Belgium), using purified oligohistidine-tagged NlpI protein for immunization. For affinity purification of the serum, purified His-NlpI (5 mg) was coupled to 0.45 g of CNBr-activated sepharose (GE) following the manufacturers protocol. The column was washed with 30 ml of wash buffer I (10 mM Tris/HCL, 1 M NaCl, 10 mM MgCl2, 0.1% Triton X-100, pH 7.2), 5 ml of elution buffer I (100 mM Glycin/HCl, 0.1% Triton X-100 pH 2.0) and equilibrated with 30 ml of block buffer (200 mM Tris/HCl, 500 mM NaCl, 0.1% Triton X-100 pH 8.0). Rabbit serum (10 ml) was mixed with 35 ml of serum buffer (10 mM Tris/HCl pH 7.4) and adjusted to a total concentration of 0.1% of Triton X-100. The solution was centrifuged (4000 × *g*, 45 min, 4°C) and the supernatant was applied to the CNBr-activated sepharose His-PBP2 column using a peristaltic pump with constant slow flow for 48 h. The column was washed with 30 ml of wash buffer I and with 20 ml of wash buffer II (10 mM Tris/HCL, 150 mM NaCl, 10 mM MgCl_2_, 0.1% Triton X-100, pH 7.2). The NlpI antibodies were eluted with 10 ml of elution buffer I and mixed with 2 ml of elution buffer II (2 M Tris/HCl, pH 8.0) afterwards. The elution was analyzed by SDS-PAGE and glycerol was added to a final concentration of 20% and the purified PBP2 antibodies were stored at −20°C. Anti-NlpI was tested for specificity by Western Blot (Figure S5C).

### Preparation of membrane fraction for affinity chromatography

This protocol was adapted from a previously published method (Vollmer *et al.*, 1999). Membranes were isolated from 4 L of *E. coli* BW25133 grown at 37°C to an optical density (578 nm) of 0.7. Cells were harvested at (5000 *× g*, 10 min, 4°C), resuspended in 20 ml of MF buffer I (10 mM Tris/maleate, 10 mM MgCl_2_, pH 6.8) and disrupted by sonication, with a Branson Digital Sonifier operating at 50 W for 5 min. Membranes were sedimented by ultracentrifugation (80000 *× g*, 60 min, 4°C). The pellet was re-suspended in 20 ml of MF buffer II (10 mM Tris-maleate, 10 mM MgCl_2_, 1 M NaCl, 2% Triton X-100, pH 6.8) to extract all membrane proteins by stirring over-night at 4°C. The supernatant obtained after another ultracentrifugation step (80000 *× g*, 60 min, 4°C), was diluted by the addition of 20 ml of MF dialysis buffer I (10 mM Tris/maleate, 10 mM MgCl_2_, 50 mM NaCl, pH 6.8) and dialyzed against 5 L of the same buffer. The obtained membrane fraction was used directly for affinity chromatography. For high salt affinity chromatography, the obtained membrane fraction was dialyzed against 3 L of MF buffer III (10 mM Tris/maleate, 10 mM MgCl_2_, 400 mM NaCl, pH 6.8). For membrane extracts using the detergent DDM, Trition X-100 was replaced with 1% DDM.

### Affinity chromatography

This protocol was adapted from a previously published method (Vollmer *et al.*, 1999). Sepharose beads were activated following the instructions of the manufacturer (GE). Coupling of 2 mg of protein to 0.13 g of activated sepharose beads was carried out overnight at 6°C with gentle agitation in protein buffer. After washing the beads with protein buffer, the remaining coupling sites were blocked by incubation in AC blocking buffer (200 mM Tris/HCl, 10 mM MgCl_2_ 500 mM NaCl, 10% glycerol and 0.25% Triton X-100, pH 7.4) with gentle agitation over-night at 6°C. The beads were washed alternatingly with AC blocking buffer and AC acetate buffer (100 mM sodium acetate, 10 mM MgCl_2_, 500 mM NaCl, 10% glycerol and 0.25% Triton X-100, pH 4.8), and finally re-suspended in AC buffer I (10 mM Tris/maleate, 10 mM MgCl_2_, 50 mM NaCl, 1% Triton X-100, pH 6.8). As control (Tris-Sepharose) one batch of activated Sepharose beads was treated identically with the exception that no protein was added. Affinity chromatography was performed at 6°C. *E. coli* membrane fraction extracted out of 2 L per sample (see above) containing 50 mM NaCl (or 400 mM NaCl for high salt chromatography) was incubated with gentle agitation over-night. The column was washed with 10 ml of AC wash buffer (10 mM Tris/maleate, 10 mM MgCl_2_, 50 mM NaCl and 0.05% Triton X-100, pH 6.8). Retained proteins were eluted with 20 ml of AC elution buffer I (10 mM Tris/maleate, 10 mM MgCl_2_, 150 mM NaCl, 0.05% Triton X-100, pH 6.8) followed by a second elution step with 1 ml of AC elution buffer II (10 mM Tris/maleate, 10 mM MgCl_2_, 1 M NaCl, 0.05% Triton X-100, pH 6.8). Both elution fractions were stored at −20°C. For the high salt affinity chromatography the AC high salt wash buffer (10 mM Tris/ maleate, 10 mM MgCl_2_, 400 mM NaCl and 0.05% Triton X-100, pH 6.8) and the AC high salt elution buffer (10 mM Tris/maleate, 10 mM MgCl_2_, 2 M NaCl, 0.05% Triton X-100, pH 6.8) were used. Elutions were analyzed by liquid chromatography (LC)-MS/MS.

### Mass spectrometry to identify NlpI affinity chromatography hits

For liquid chromatography (LC)-MS/MS, tryptic peptides were desalted (Oasis HLB μElution Plate, Waters), dried in vacuum and reconstituted in 20 μl of 4% acetonitrile, 0.1% formic acid. In total 1 μg of peptide was separated with a nanoACQUITY UPLC system (Waters) fitted with a trapping column (nanoAcquity Symmetry C_18_; 5 μm [average particle diameter]; 180 μm [inner diameter] × 20 mm [length]) and an analytical column (nanoAcquity BEH C_18_; 1.7 μm [average particle diameter]; 75 μm [inner diameter] × 200 mm [length]). Peptides were separated on a 240 min gradient and were analyzed by electrospray ionization–tandem mass spectrometry on an Orbitrap Velos Pro (Thermo Fisher Scientific). Full-scan spectra from a mass/charge ratio of 300 to one of 1,700 at a resolution of 30,000 full width at half maximum were acquired in the Orbitrap mass spectrometer. From each full-scan spectrum, the 15 ions with the highest intensity were selected for fragmentation in the ion trap. A lock-mass correction with a background ion (mass/charge ratio, 445.12003) was applied. The raw mass spectrometry data was processed with MaxQuant (v1.5.2.8) (Cox & Mann, 2008) and searched against an Uniprot *E.coli* K12 proteome database. The search parameters were as following: Carbamidomethyl (C) (fixed), Acetyl (N-term) and Oxidation (M) (variable) were used as modifications. For the full scan MS spectra (MS1) the mass error tolerance was set to 20 ppm and for the MS/MS spectra (MS2) to 0.5 Da. Trypsin was selected as protease with a maximum of two missed cleavages. For protein identification a minimum of one unique peptide with a peptide length of at least seven amino acids and a false discovery rate below 0.01 were required on the peptide and protein level. The match between runs function was enabled, a time window of one minute was set. Label free quantification was selected using iBAQ (calculated as the sum of the intensities of the identified peptides and divided by the number of observable peptides of a protein) (Schwanhausser *et al.*, 2011), with the log fit function enabled.

The proteinGroups.txt file, an output of MaxQuant, was loaded into R (ISBN 3-900051-07-0) for further analysis. The iBAQ-values of the MaxQuant output were first batch-corrected using the limma package (Ritchie *et al.*, 2015) and then normalized with the vsn package (Huber *et al.*, 2002). Individual normalization coefficients were estimated for each biological condition separately. Limma was used again to test the normalized data for differential expression. Proteins were classified as a ‘hit’ with a log2 fold change higher than 4 and a ‘candidate’ with a log2 fold change higher than 2.

Data availability: The mass spectrometry proteomics data will be deposited to the ProteomeXchange Consortium via the PRIDE partner repository during the publication process.

### Genetic interaction assay

For quantitation of genetic interactions; strains were grown to late exponential phase (~0.7 OD_578_), adjusted to an OD_578_ of 1 and spread out using glass beads on rectangular LB Lennox plates (200 μl per strain per plate). Plates were dried at 37°C for one hour and arrayed using a Rotor HDA replicator on Lennox LB agar plates. On each genetic interaction assay plate the parental strain, the single deletion A, the single deletion B and the double deletion AB (or BA) were arrayed, each in 96 copies per plate. Plates were incubated at 37°C for 12 h and imaged under controlled lighting conditions (spImager S&P Robotics) using an 18 megapixel Canon Rebel T3i (Canon). Colony integral opacity as fitness readout was quantified using the image analysis software Iris (Kritikos *et al.*, 2017). Double mutant genetic interaction scores were calculated as previously described. Briefly, fitness ratios are calculated for all mutants by dividing their fitness values by the respective WT fitness value. The product of single mutant fitness ratios (expected) is compared to the double mutant fitness ratio (observed) across replicates. The probability that the two means (expected and observed) are equal across replicates is obtained by a Student’s two-sample t-test.

### Thermal proteome profiling and sample preparation

Thermal proteome profiling was performed as previously described in (Mateus *et al.*, 2018). Briefly, bacterial cells were grown overnight at 37°C in lysogeny broth, and diluted 100-fold into 20 ml of fresh medium. Cultures were grown aerobically at 37°C with shaking until optical density at 578 nm (OD_578_) ~0.5. Cells were then pelleted at 4000 × *g* for 5 min, washed with 10 ml PBS, re-suspended in the same buffer to an OD_578_ of 10, and aliquoted to a PCR plate. The plate was subjected to a temperature gradient for 3 minutes in a PCR machine (Agilent SureCycler 8800), followed by 3 minutes at room temperature. Cells were lysed with lysis buffer (final concentration: 50 µg/ml lysozyme, 0.8% NP40, 1X protease inhibitor (Roche), 250 U/ml benzonase and 1 mM MgCl_2_ in PBS) for 20 min, shaking at room temperature, followed by three freeze-thaw cycles. Protein aggregates were then removed and the soluble fraction was digested according to a modified SP3 protocol (Mateus *et al.*, 2018). Peptides were labelled with TMT6plex (ThermoFisher Scientific), desalted with solid-phase extraction on a Waters OASIS HLB µElution Plate (30 µm), and fractionated onto six fractions on a reversed phase C18 system running under high pH conditions.

### 2D-TPP mass spectrometry-based proteomics

Samples were analyzed with liquid chromatography coupled to tandem mass spectrometry, as previously described (Mateus *et al.*, 2018). Briefly, peptides were separated using an UltiMate 3000 RSLC nano LC system (Thermo Fisher Scientific) equipped with a trapping cartridge (Precolumn C18 PepMap 100, 5µm, 300 µm i.d. x 5 mm, 100 Å) and an analytical column (Waters nanoEase HSS C18 T3, 75 µm x 25 cm, 1.8 µm, 100 Å). Solvent A was 0.1% formic acid in LC-MS grade water and solvent B was 0.1% formic acid in LC-MS grade acetonitrile. After loading the peptides onto the trapping cartridge (30 µL/min of solvent A for 3 min), elution was performed with a constant flow of 0.3 µL/min using a 60 to 120 min analysis time (with a 2–28%B elution, followed by an increase to 40%B, and re-equilibration to initial conditions). The LC system was directly coupled to a Q Exactive Plus mass spectrometer (Thermo Fisher Scientific) using a Nanospray-Flex ion source and a Pico-Tip Emitter 360 µm OD x 20 µm ID; 10 µm tip (New Objective). The mass spectrometer was operated in positive ion mode with a spray voltage of 2.3 kV and capillary temperature of 320°C. Full scan MS spectra with a mass range of 375–1200 m/z were acquired in profile mode using a resolution of 70,000 (maximum fill time of 250 ms or a maximum of 3e6 ions (automatic gain control, AGC)). Fragmentation was triggered for the top 10 peaks with charge 2 to 4 on the MS scan (data-dependent acquisition) with a 30 s dynamic exclusion window (normalized collision energy was 32), and MS/MS spectra were acquired in profile mode with a resolution of 35,000 (maximum fill time of 120 ms or an AGC target of 2e5 ions).

### 2D-TT data analysis

Protein identification and quantification. Mass spectrometry data were processed as previously described (Mateus *et al.*, 2018). Briefly, raw mass spectrometry files were processed with IsobarQuant (Franken *et al.*, 2015) and peptide and protein identification were performed with Mascot 2.4 (Matrix Science) against the *E. coli* (strain K12) Uniprot FASTA (Proteome ID: UP000000625), modified to include known contaminants and the reversed protein sequences (search parameters: trypsin; missed cleavages 3; peptide tolerance 10ppm; MS/MS tolerance 0.02Da; fixed modifications were carbamidomethyl on cysteines and TMT10plex on lysine; variable modifications included acetylation on protein N-terminus, oxidation of methionine and TMT10plex on peptide N-termini).

Thermal proteome profiling analysis. Data analysis was performed in R, as previously described in (Mateus *et al.*, 2018). Briefly, all output data from IsobarQuant was normalized using variance stabilization (*vsn*) (Huber *et al.*, 2002). Abundance and stability scores were calculated with a bootstrap algorithm (Becher *et al.*, 2018), together with a local FDR that describes the quality and the reproducibility of the score values (by taking into account the variance between replicates). A local FDR<0.05 and a minimum absolute score of 10 were set as thresholds for significance. Abundance and stability scores of knocked out genes were discarded.

Data availability: The mass spectrometry proteomics data will be deposited to the ProteomeXchange Consortium via the PRIDE partner repository during the publication process.

**Table 1.**
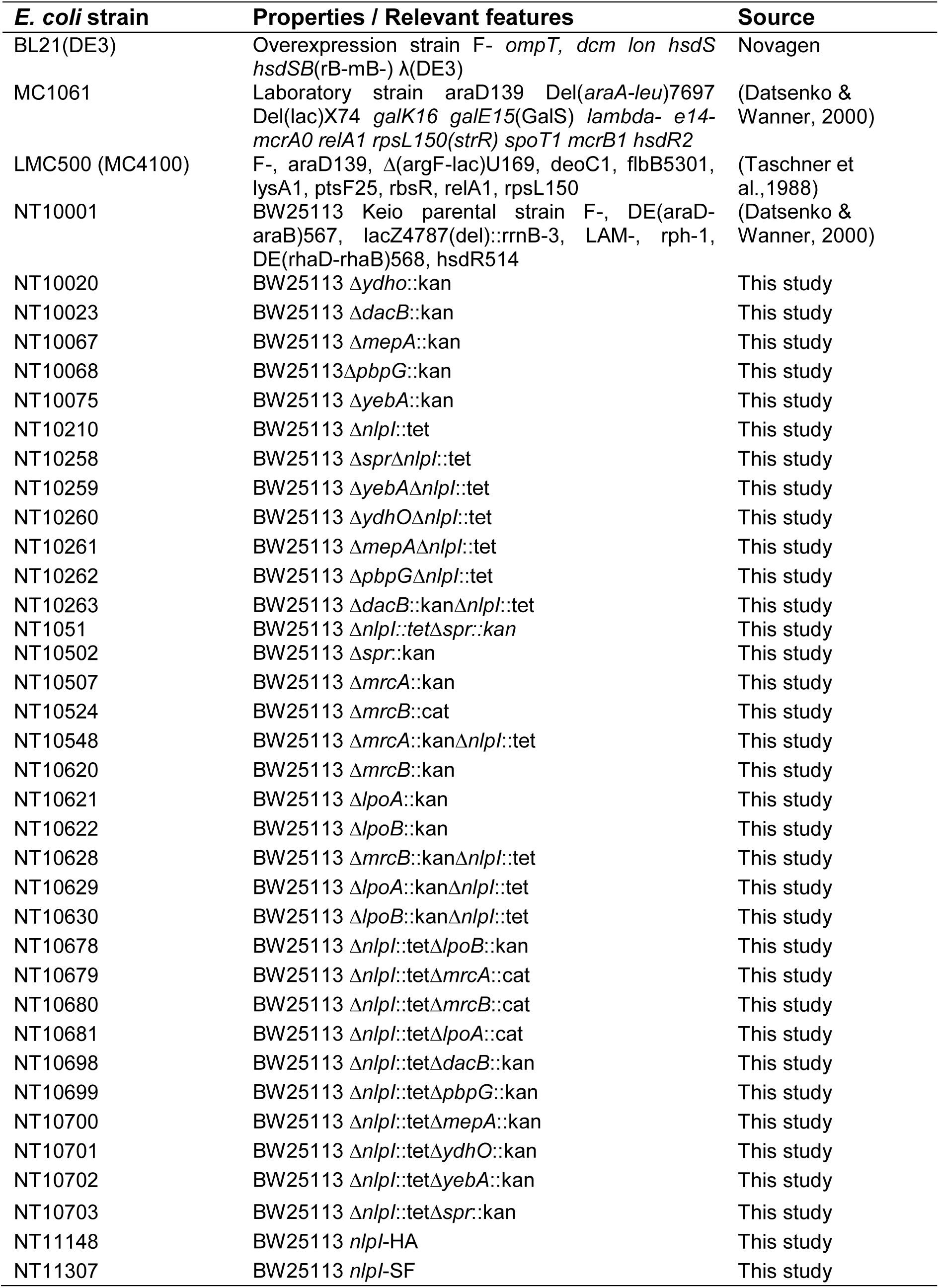
Strain list.

**Table 2.**
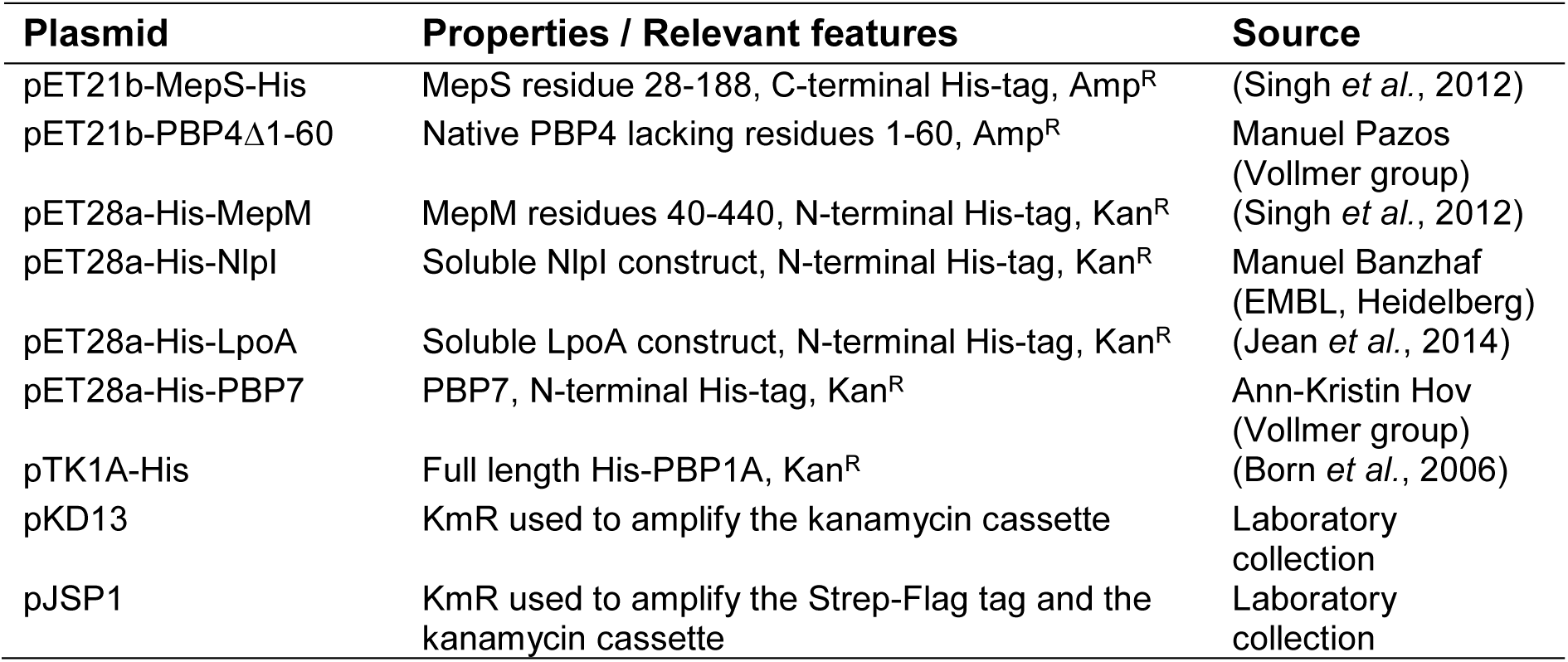
Plasmid list.

**Table 3.**
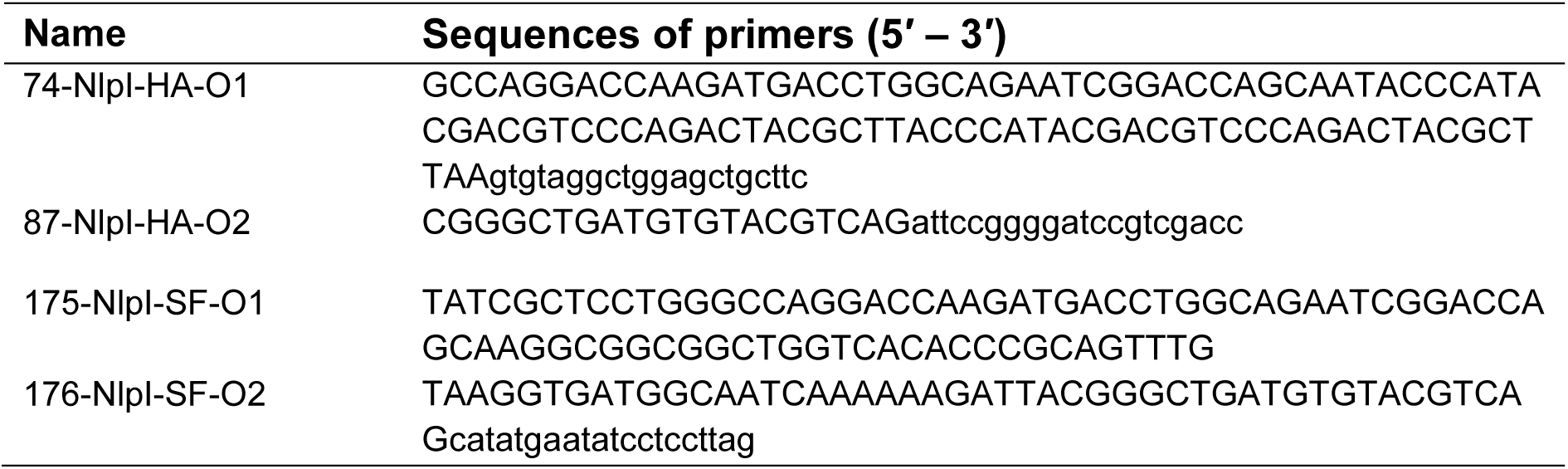
Primer list.

## Acknowledgements

We want to thank Manjula Reddy for fruitful discussions. This work was supported by the Sofja Kovalevskaja Award of the Alexander von Humboldt Foundation and EMBL core funding to A.T., the Wellcome Trust (101824/Z/13/Z) and Medical Research Council (MR/N002679/1) to V.W.. M.B. is funded by the Royal Society (RGS\R1\191041). A.M. is supported by a fellowship from the EMBL Interdisciplinary Postdoc (EI3POD) programme under Marie Skłodowska-Curie Actions COFUND (grant number 664726). M.W. was supported by a Humboldt postdoctoral fellowship.

## Author contributions

M.B, H.C.L.Y and J.V; acquisition, analysis and interpretation of data, drafting and revising article. A.L, G.K, A.M, A-K.H, F.S, M.W, M.P, A.S and M.S; acquisition of data, analysis and interpretation of data, revising the article. T.dB, A.T and W.V; conception and design, analysis and interpretation of data, drafting and revising the article.

## SUPPLEMENTARY INFORMATION

**Supplementary table 1:** 2DTPP raw data

**Supplementary table 2:** 2DTPP hits

**Supplementary table 3:** AP-MS-all raw data

**Supplementary table 4:** AP-MS-hits

**Supplementary table 5:** fitness.ratios.csv

**Fig. S1.**
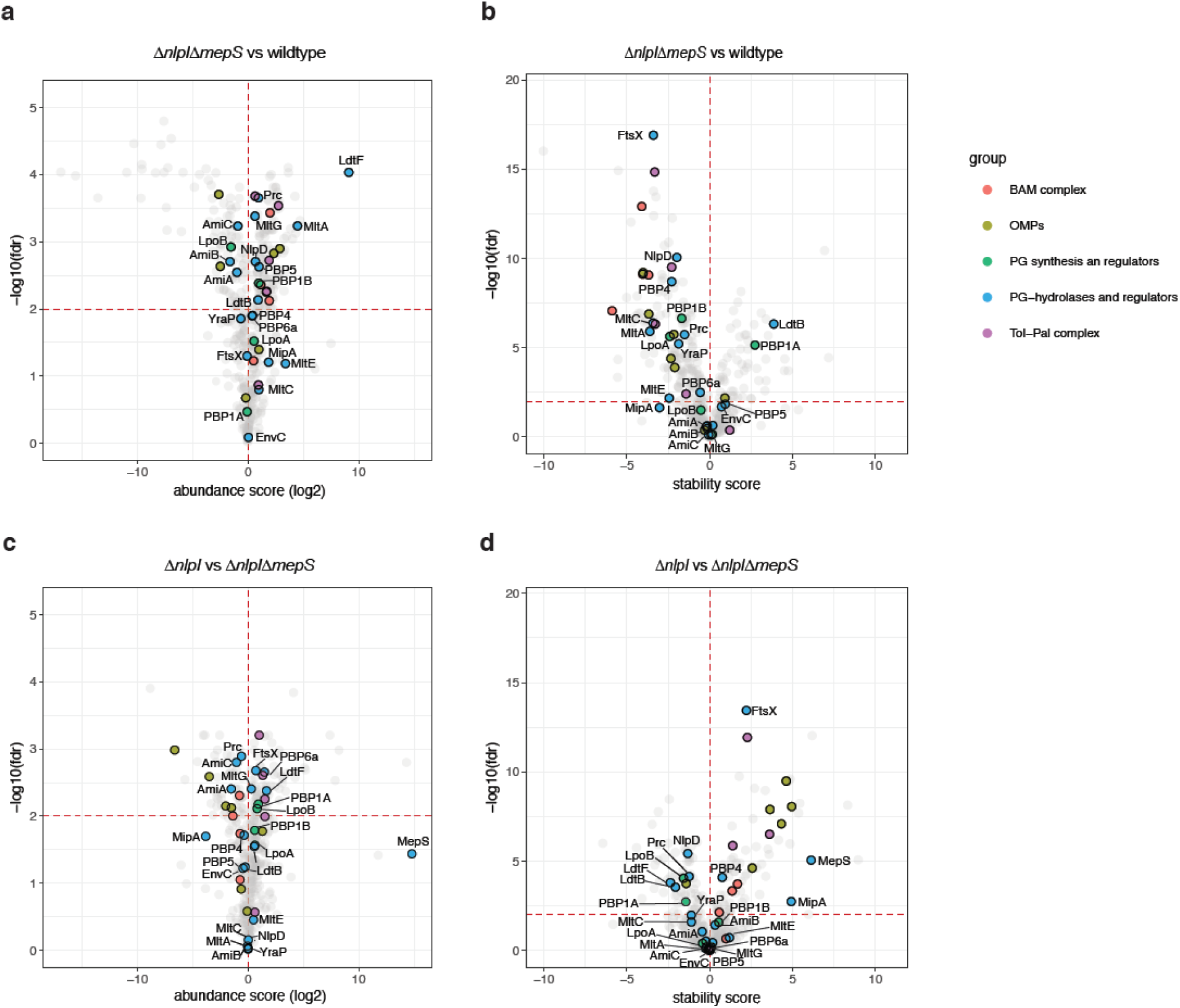
*In vivo* and *in vitro* proteomic interaction assays link NlpI to several classes of PG hydrolases. **a-b.** Wild-type and Δ*nlpI*Δ*mepS* cells were heated at a range of temperatures and the soluble components were labelled, combined and quantified by LC-MS, using the published 2D-TPP protocol (Mateus *et al.*, 2018). Here shown volcano plots of two replicates: protein abundance (a) and thermostability (b). A local FDR<0.01 was set as a threshold for significance. Abundance and stability scores of knocked out genes were discarded. Highlighted: outer membrane porins (OMPs, light green), β-barrel assembly machinery (BAMs, red), PG synthases and regulators (green), PG hydrolases and regulators (blue) and the Tol-Pal complex (violet). Full results can be found in supplementary tables 1 and 2. **c-d.** 2D-TPP profiles of Δ*nlpI* compared to Δ*nlpI*Δ*mepS* cells. Shown in the volcano plots are changes in protein abundance (c) and thermostability (d). A local FDR<0.05 and a minimum absolute score of 10 were set as thresholds for significance. Highlighted: outer membrane porins (OMPs, light green), β-barrel assembly machinery (BAMs, red), PG synthases and regulators (green), PG hydrolases and regulators (blue) and Tol-Pal complex proteins (violet). Full results can be found in supplementary tables 1 and 2.

**Fig. S2.**
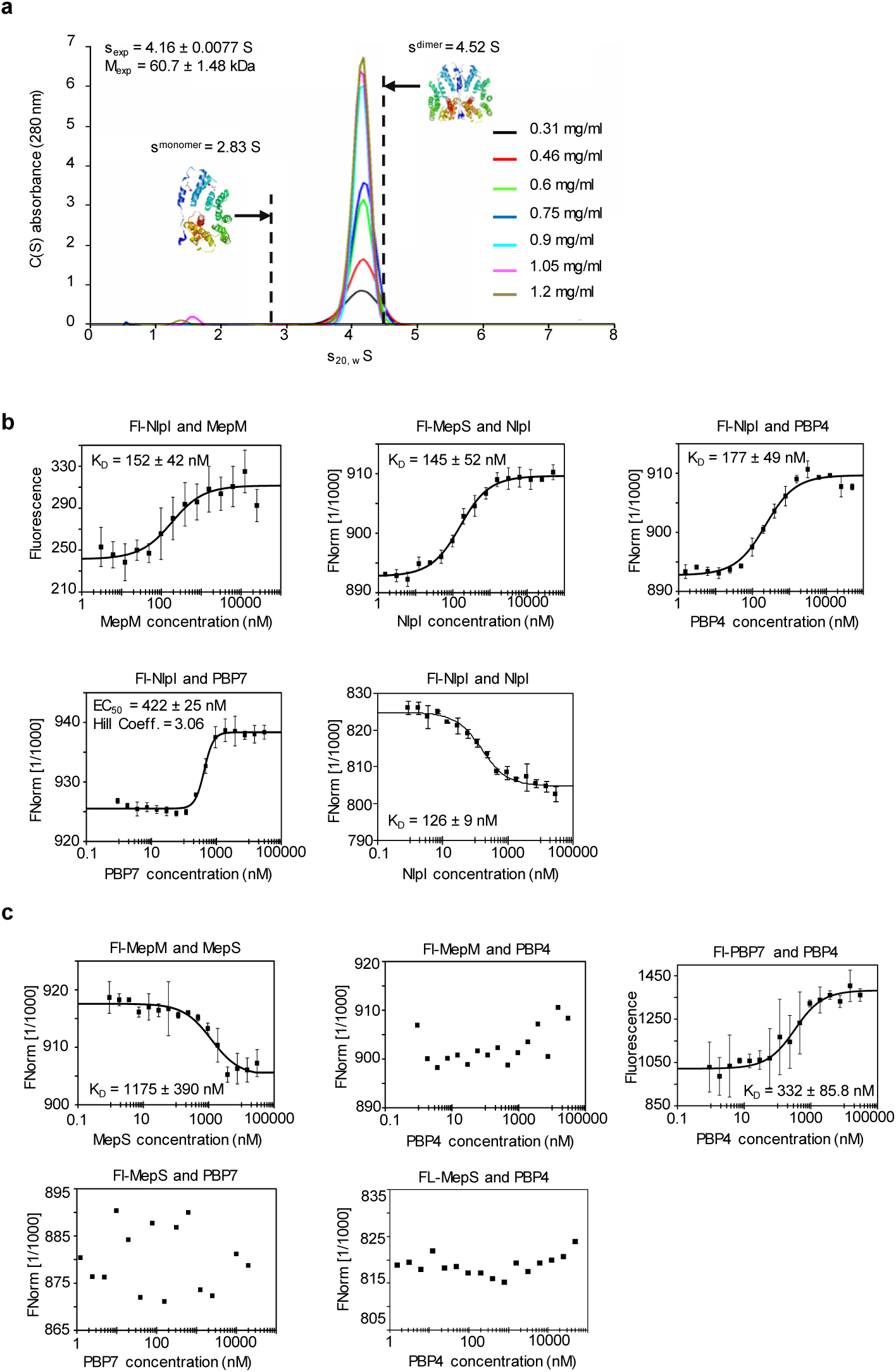
NlpI dimerizes and interacts with several endopeptidases *in vitro*. **a.** Dimer of NlpI detected by Analytical ultracentrifugation (AUC). The soluble version of NlpI (residues 19-294) has a molecular weight of 30.35 kDa. **b.** Microscale thermophoresis (MST) binding curves for interactions between fluorescently labelled MepS (fl-MepS) with unlabelled NlpI and fl-NlpI with unlabelled NlpI, MepM, PBP4 and PBP7. Ligand binding can affect thermophoresis differently; hence the binding of a ligand can enhance or reduce thermophoresis. This results in either a higher or lower FNorm in the bound state compared to the unbound state (Jerabek-Willemsen *et al.*, 2011). **c.** Binding curves for MST assays between fl-MepM with MepS and PBP4, fl-PBP7 with PBP4 and fl-MepS with PBP4 and PBP7.

**Fig S3.**
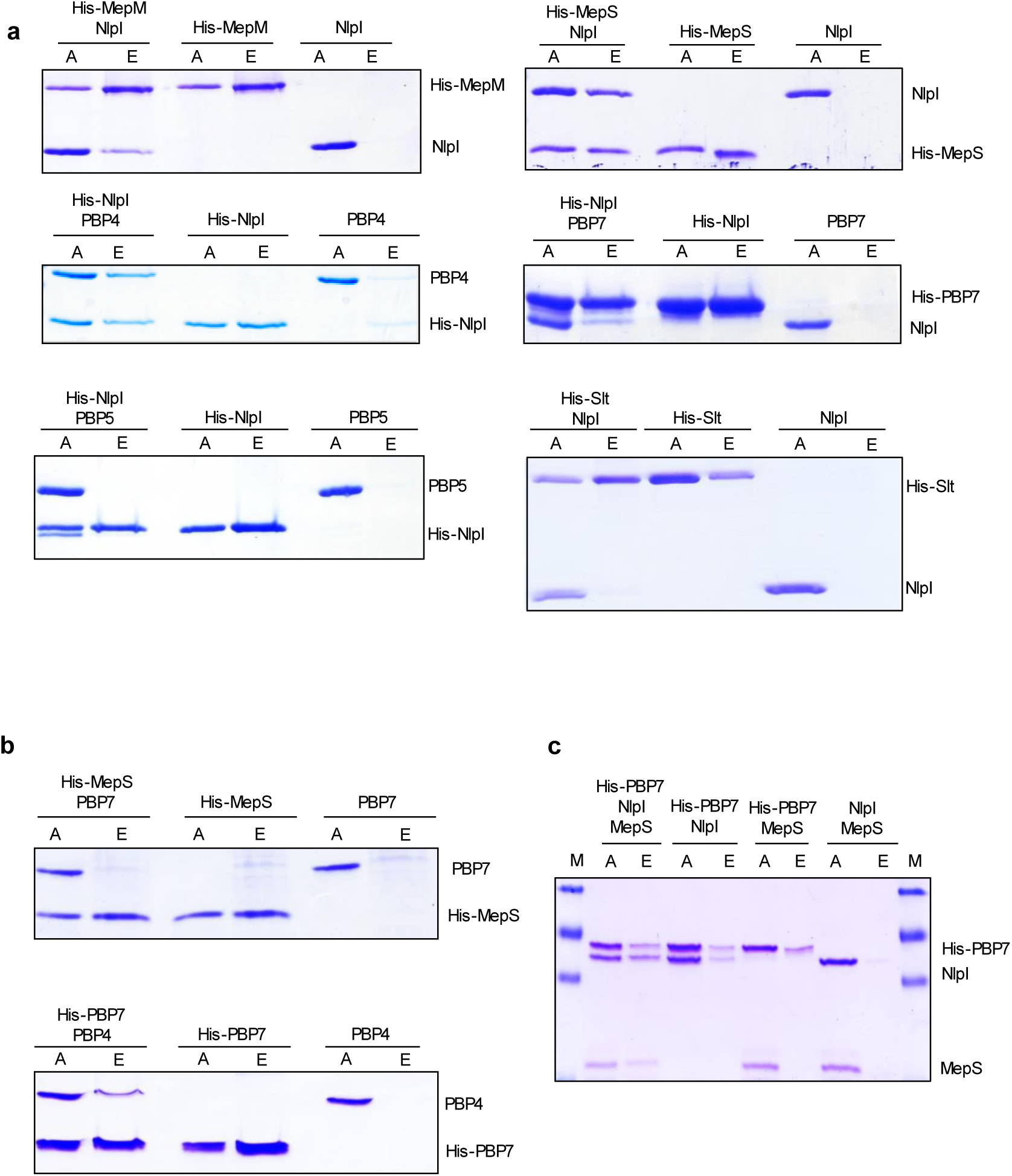
Assaying for interactions between endopeptidases and with NlpI by Ni^2+^-NTA pulldown assay. **a.** SDS-PAGE (12-15%) analysis of applied (A) and eluted (E) samples from Ni^2+^-NTA pull down assay of NlpI-EPase combinations. Equimolar concentrations of proteins for each respective assay (1 or 2 μM) were incubated with Ni^2+^-NTA beads in combination or alone. Retention of tagless protein in the presence of His-tagged partner indicates an interaction. Tagless PBP5 is not retained in the presence of His-tagged NlpI nor is tagless NlpI retained by His-Slt. **b.** SDS-PAGE (12-15%) analysis of applied (A) and eluted (E) samples from Ni^2+^-NTA pull down assay of His-MepS with PBP7 and His-PBP7 with PBP4. Equimolar concentrations of proteins for each respective assay (1-2 μM) were incubated with Ni^2+^-NTA beads in combination or alone. **c.** SDS (15%) analysis of applied (A) and eluted (E) samples from Ni^2+^-NTA pull down assay of His-PBP7 with NlpI (both 2 µM) and MepS (4 µM) in the presence of formaldehyde crosslinker. M, Marker.

**Fig. S4.**
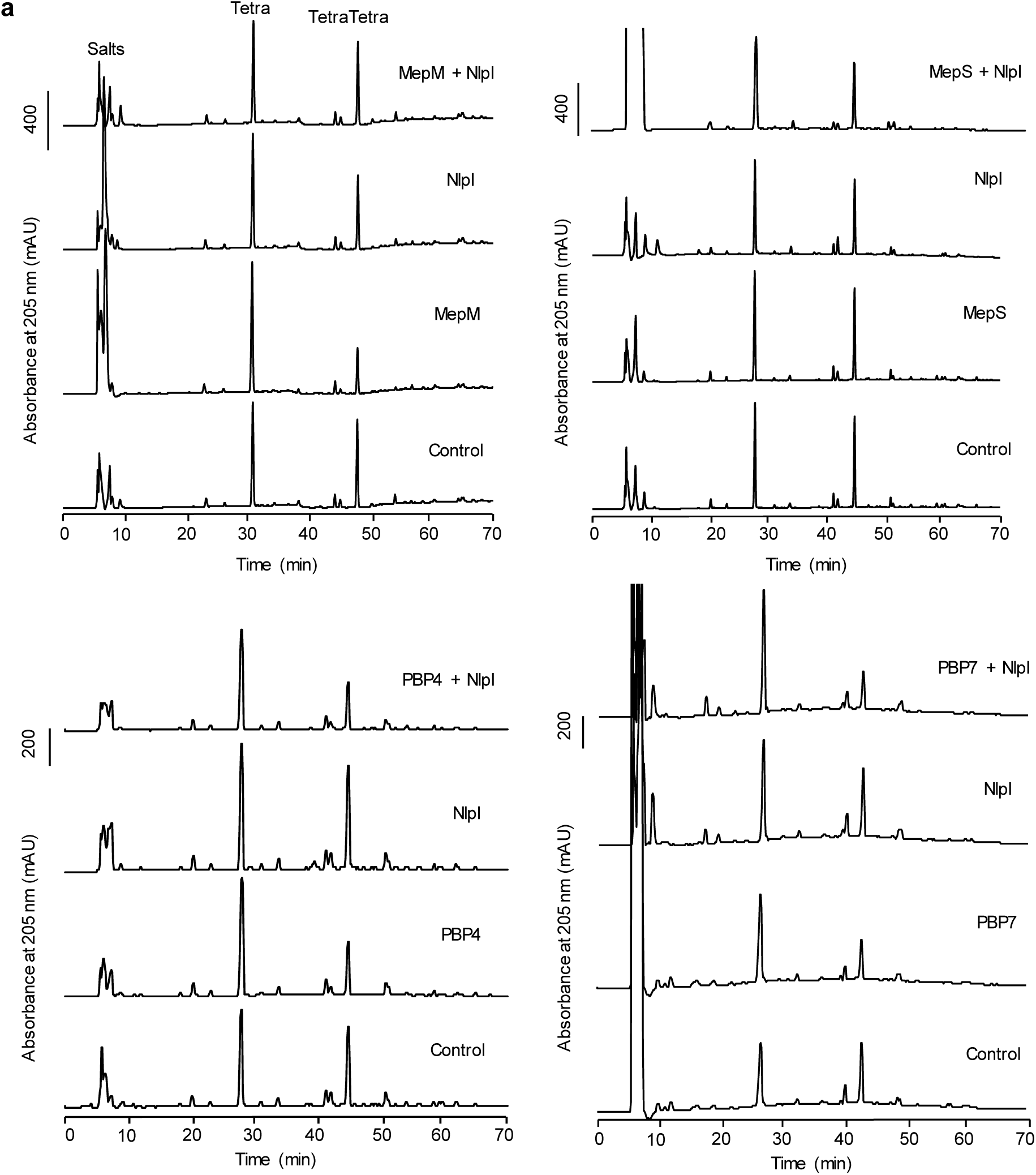
*in vitro* PG digestion assays. NlpI moderately affects the activity of MepM, but less so the activity of MepS, PBP4 and PBP7. Representative HPLC chromatograms of assays containing the respective EPase with or without NlpI. Control samples were incubated with no enzyme. Muropeptides were separated and the PG profiles determined as previously described (Glauner, 1988). Figure shows representative chromatograms of 3-6 independent experiments.

**Fig. S5.**
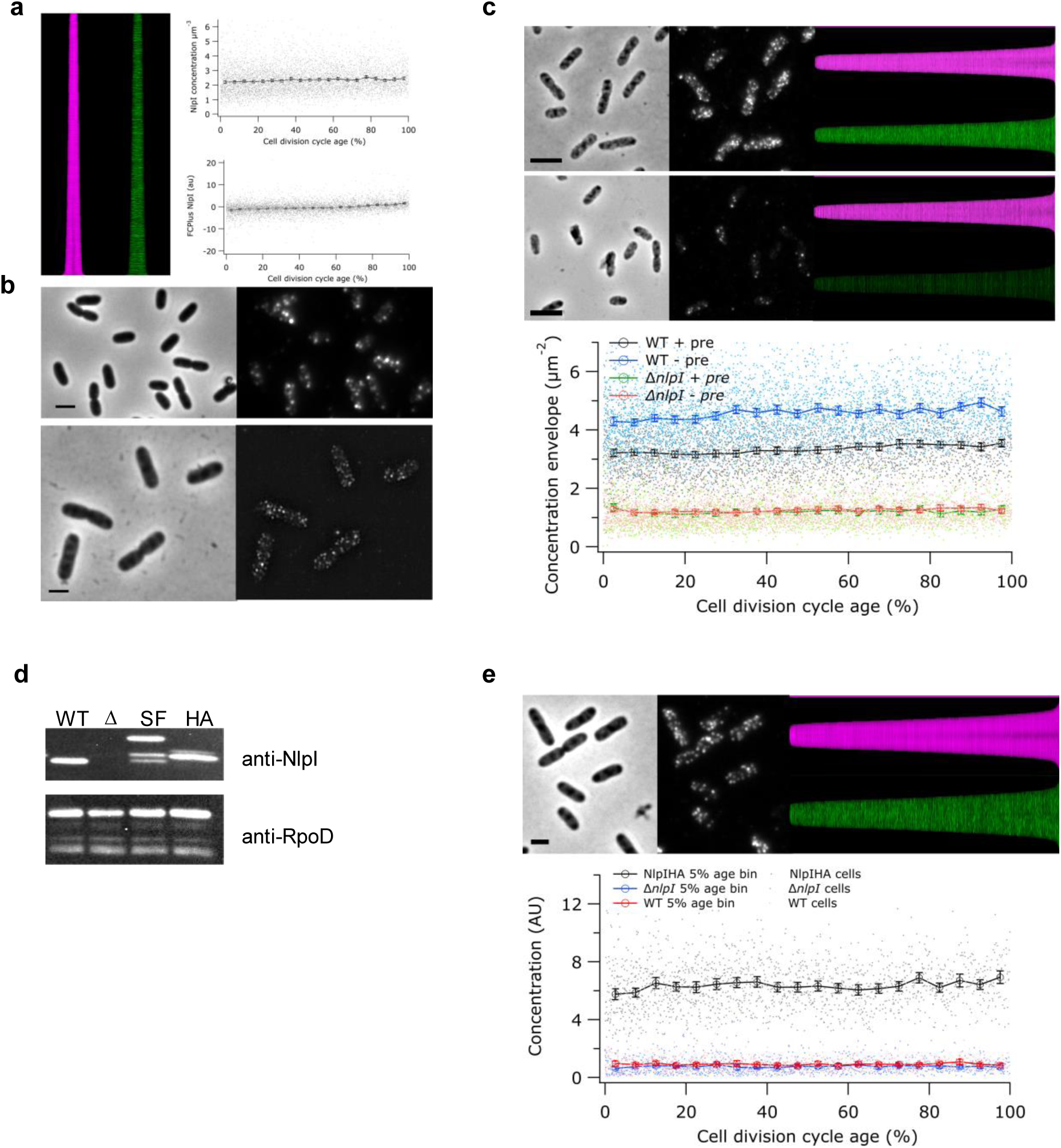
NlpI localizes along the cell envelope. **a.** MC4100 cells grown to steady state in minimal glucose medium at 28°C were immunolabeled with antibodies specific for NlpI. The map of diameters (magenta) and map of fluorescence (green) of NlpI localization where cells are sorted according to their cell length are shown. The concentration of NlpI in molecules per µm^−2^ wall area as function of the normalized cell division cycle age is constant. The grey dots are the values measured for the individual cells and the black markers are the 5% age bins with the 95 confidence error bars. The extra fluorescence at midcell compared to the rest of the cell (FCPlus) plotted as function of the normalized cell division cycle. NlpI is not specifically present at midcell (number of analyzed cells is 6087). **b.** A high resolution fluorescence SIM image of BW23115 cells that were grown in LB at 37°C immunolabeled with anti-NlpI on the right and the corresponding Phase contrast image on the left is shown in the bottom panel. Scale bar equals 2 µm. **c.** Wildtype BW25113 and the isogenic Δ*nlpI* strain were grown in TY at 37°C and fixed. The serum against NlpI was absorbed (pre) to the Δ*nlpI* cells and the remaining antibodies were used to label the wildtype strain (upper images) and another sample of the Δ*nlpI* cells (lower images). From left to right are shown: a phase contrast and the corresponding fluorescence image of the labelled cells and the map of diameters (magenta) and map of fluorescence (green) of the cells sorted according to ascending length. Scale bar equals 2µm. Graph of the concentration NlpI (AU) in the envelope of the cells plotted against the cell division cycle time (%) are also shown. The dots are the data on the individual cells, whereas the markers are the 5% age bins with the 95 confidence error bars. **d**. Western blot using anti-NlpI for constructs carrying different chromosomally tagged NlpI versions, and the wild-type and Δ*nlpI* controls. Cells were grown exponentially for 2 hours, harvested and adjusted to 2 μg total protein per lane. Samples were separated by SDS PAGE and transferred by Western Blot on a PVDF membrane. NlpI was visualized by incubation with anti-NlpI followed by incubation with anti-rabbit HRP. Samples are as followed: WT: BW25113; Δ: BW25113Δ*nlpI*::*kan*; SF: BW25113*nlpI*::*strep*::*flag*::*kan*; HA: BW25113*nlpI*::*HA*::*frt*. Anti-NlpI is able to visualise NlpI. **e.** Wildtype BW25113 and the isogenic Δ*nlpI* strain and the same strain chromosomally expressing NlpI-HA were grown in TY at 37°C and fixed and labelled with anti HA. From left to right are shown: a phase contrast and the corresponding fluorescence image of the labelled cells and the map of diameters (magenta) and map of fluorescence (green) of the cells sorted with ascending length. Scale bar equals 1 µm. Also shown in panel D is a graph of the concentration HA-NlpI (AU) in the cells plotted against the cell division cycle time (%). The dots are the data on the individual cells, whereas the markers are the 5% age bins with the 95% confidence error bars.

**Fig. S6.**
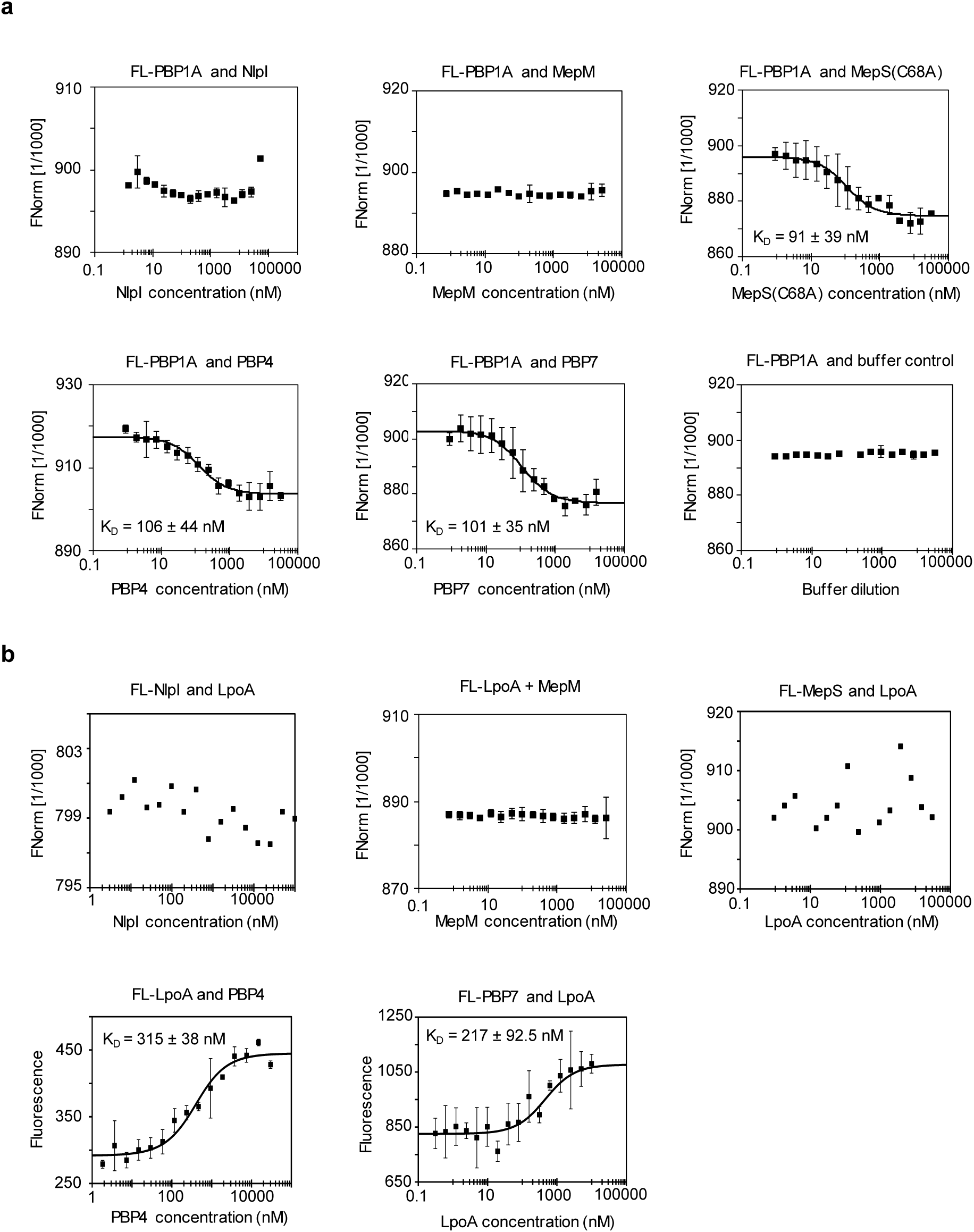
Reconstitution of PG multi-enzyme complexes with PBP1A and LpoA. **a.** MST curves assaying for interaction between PBP1A with NlpI, MepM, MepS(C68A), PBP4, PBP7 and buffer control. A catalytically inactive version of MepS (MepS(C68A)) was used as unlabelled ligand in these experiments. **b.** MST curves assaying for interaction between LpoA with NlpI, MepM, MepS, PBP4, PBP7.

**Fig. S7.**
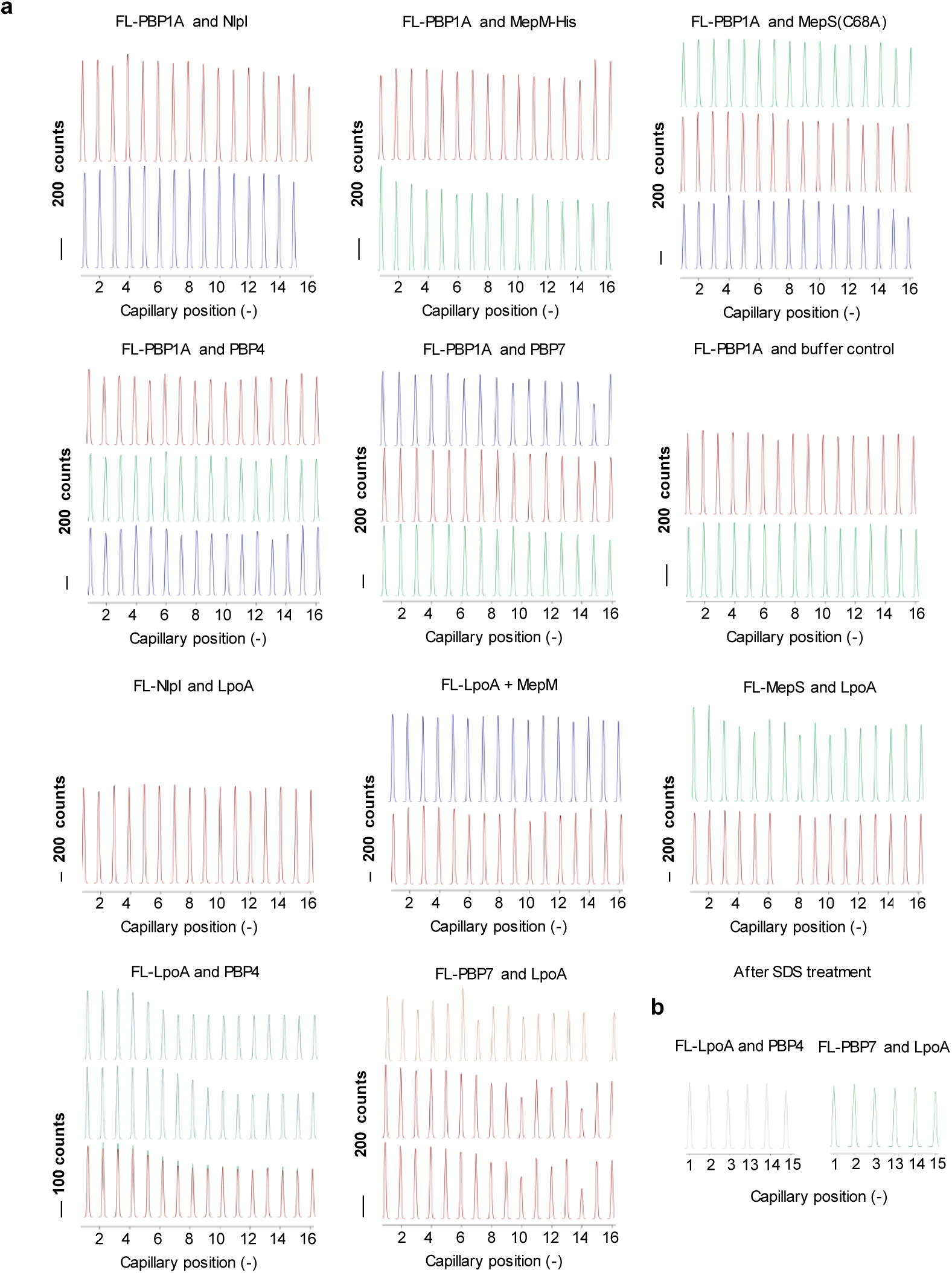
Initial fluorescence scans for MST experiments between PBP1A/LpoA with NlpI/EPases. **a.** MST capillary scans showing initial fluorescence of respective reactions testing for interactions between PBP1A/LpoA against NlpI and EPases. Scale bar = 200 fluorescence counts. **b.** Capillary scans for samples showing ligand concentration dependent changes in fluorescence were repeated after boiling samples in reducing agent and SDS to abolish ligand binding. The fluorescence reads for samples containing highest and lowest concentration of ligand were within the margin of error, suggesting that differences in initial fluorescence were due to ligand binding and not different Fl-protein concentration, validating the use of the raw fluorescence data to plot binding curves.

**Fig. S8.**
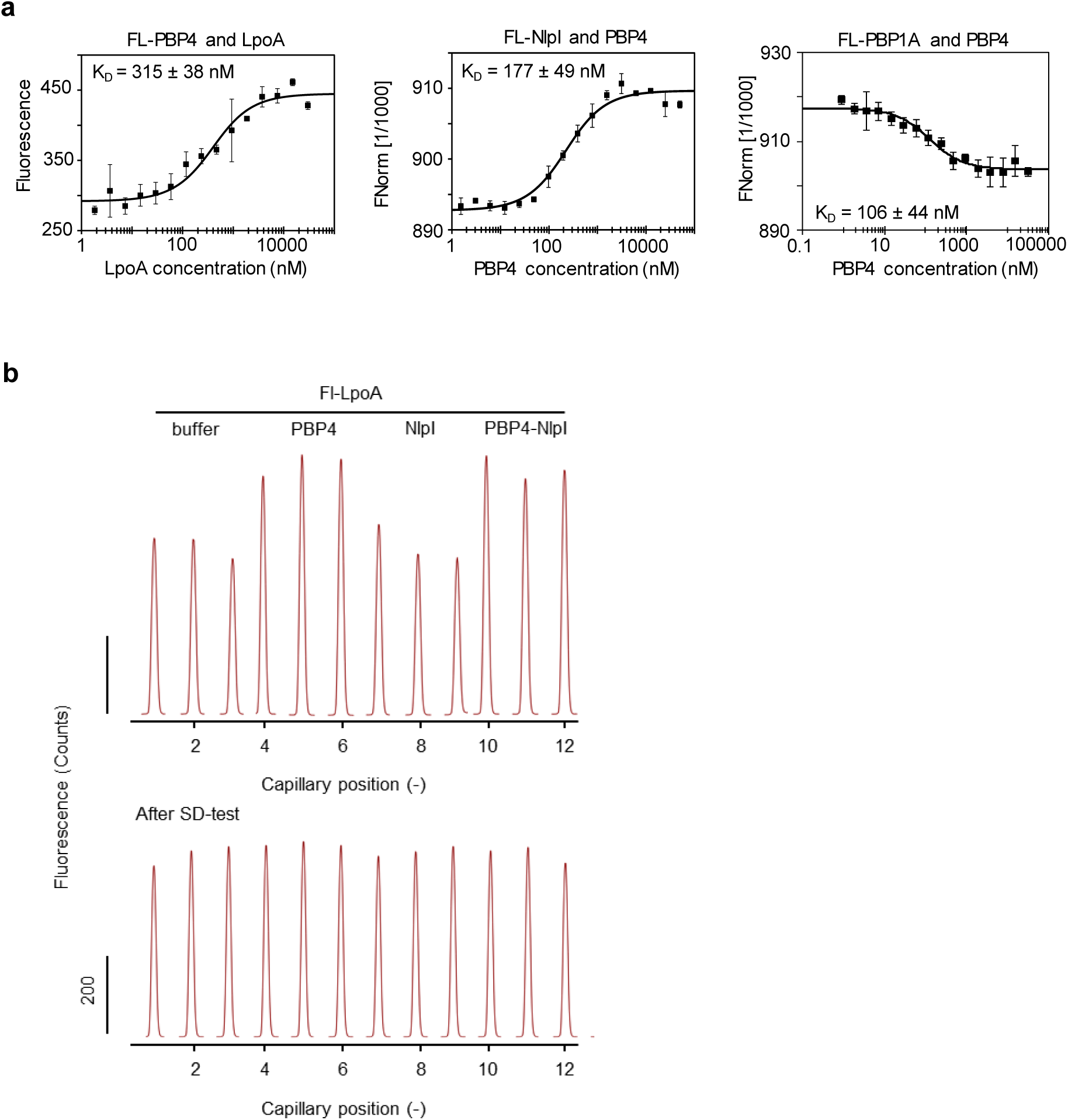
PBP4 interacts simultaneously with NlpI and the PBP1A-LpoA synthase complex. **a.** MST interaction curves of PBP4 with PBP1A, LpoA and NlpI, respectively. **b.** Fixed concentration MST assays with LpoA-PBP4-NlpI showed differences in initial fluorescence. These differences were eliminated upon boiling of samples in SDS (which abolishes ligand binding), which suggests that the initial differences were due to ligand binding and not due to inaccurate pipetting, validating the use of the raw fluorescence data to plot binding curves.

